# Multiomics study of *CHCHD10^S59L^*-related disease reveals energy metabolism downregulation: OXPHOS and β-oxidation deficiencies associated with lipids alterations

**DOI:** 10.1101/2023.01.19.524672

**Authors:** Blandine Madji Hounoum, Rachel Bellon, Emmanuelle C Genin, Sylvie Bannwarth, Antoine Lefevre, Lucile Fleuriol, Delphine Debayle, Anne-Sophie Gay, Agnès Petit-Paitel, Sandra Lacas-Gervais, Hélène Blasco, Patrick Emond, Veronique Paquis-Flucklinger, Jean-Ehrland Ricci

## Abstract

Mutations in the coiled-coil-helix-coiled-coil-helix domain containing 10 (*CHCHD10*) gene have been associated with a large clinical spectrum including myopathy, cardiomyopathy and amyotrophic lateral sclerosis (ALS). Herein, we analyzed the metabolic changes induced by the p.S59L *CHCHD10* mutation to identify new therapeutic opportunities. Using metabolomic, lipidomic and proteomic analysis we observed a strong alteration of metabolism in plasma and heart of *Chchd10*^*S59L/+*^ mice compared to their wild type littermates at pre-symptomatic and symptomatic stages. In plasma, levels of phospholipids were decreased while those of carnitine derivatives and most of amino acids were increased. The cardiac tissue from *Chchd10*^*S59L/+*^ mice showed a decreased Oxidative Phosphorylation (OXPHOS) and β-oxidation proteins levels as well as tricarboxylic acid cycle (TCA) intermediates and carnitine pathway metabolism. In parallel, lipidomics analysis reveals a drastic change in the lipidome, including triglycerides, cardiolipin and phospholipids. Consistent with this energetic deficiency in cardiac tissue, we show that L-acetylcarnitine supplementation improves the mitochondrial network length in IPS-derived cardiomyocytes from a patient carrying the *CHCHD10*^*S59L/+*^ mutation. These data indicate that a bioenergetic intermediate such as L-acetylcarnitine may restore mitochondrial function in *CHCHD10*-related disease, due to the reduction in energy deficit that could be compensated by carnitine metabolic pathways.

## Introduction

Coiled-coil-helix-coiled-coil-helix domain containing 10 (CHCHD10) protein is localized in the intermembrane space of mitochondria, enriched at cristae junctions and involved in the stability of the mitochondrial contact site and cristae organizing system (MICOS) [1, 2]. In mitochondria, CHCHD10 dimerizes with its paralog CHCHD2 in associated complexes [3, 4]. In mice, CHCHD10 is expressed in many tissues, especially in energy-demanding tissues such as heart and muscle [3]. *In vitro* and *in vivo* studies have shown that *CHCHD10* mutants cause disease by a toxic gain-of-function mechanism rather than by loss of function [3, 5]. However, the mechanism of pathogenesis of *CHCHD10* mutations remains unclear and its function is not fully understood. We recently identified *CHCHD10* as the first genetic evidence demonstrating the primary role of mitochondria in some motor neuron disease [2, 6]. We generated knock-in (KI) mice (*Chchd10*^*S59L/+*^) that develop Amyotrophic Lateral Sclerosis (ALS) phenotype, myopathy and cardiomyopathy associated with morphological abnormalities of mitochondria, severe oxidative phosphorylation (OXPHOS) deficiency and multiple defects of respiratory chain (RC) complexes activity in several tissues in the late stage [7]. Similar results were obtained by Anderson and collaborators [5]. The discovery of a causative mutation in the mitochondrial protein CHCHD10 provided direct evidence that disruption of mitochondrial structure may contribute to the etiology of ALS. It has been described that ALS is also associated with altered energy metabolism, including weight loss, hypermetabolism and hyperlipidaemia [8-11]. Furthermore, regarding the crucial role of mitochondria function in cardiac health, mitochondrial cardiomyopathies are known to be associated with metabolic remodeling [12, 13]. Thus, we hypothesize that metabolic disturbances are key pathogenic mechanisms related to mitochondrial defects in *CHCHD10*-related diseases.

In this study, we aim to understand how mitochondria dysfunction associated with *CHCHD10* mutations triggers altered global metabolism. We used “multiomics” approaches including metabolomics, lipidomics and proteomics analysis of plasma and heart from *Chchd10*^*S59L/+*^ mice and their wild-type littermates. We observed a significant metabolic change in *Chchd10*^*S59L/+*^ heart with downregulation of energy metabolism intermediates, OXPHOS and β-oxidation protein levels, and a dramatic change in the lipidome. Furthermore, using IPS-derived cardiomyocytes from patient carrying the CHCHD10^S59L/+^ mutation, we found that mitochondrial dysfunction could be reversed by supplementation with bioenergy intermediates such as L-acetylcarnitine.

## Materials and Methods

### Animals

All animal procedures and experiments were approved by the local ethical committee (Comité d’Ethique en Expérimentation Animale IGBMC-ICS) for Animal Care and Use and MESR. *Chchd10*^*S59L/+*^ mice and their wild-type littermate were the same as from the previous study (Genin et al., 2019). The phenotypical characteristics and biochemistry measurement of the mice were performed at Phenomin-ICS, Illkirch, France. For these studies, the population included 40 C57BL/6N mice (20 males and 20 females) divided in 2 groups (10 *Chchd10*^*S59L/+*^ and 10 wild-type of each sex).

For metabolomics and lipidomics analysis, cardiac tissue was collected from symptomatic *Chchd10*^*S59L/+*^ mice (40 to 46 weeks of ages) and their wild-type littermate (n=12 mice per group). No statistical difference was found regarding age and gender between the symptomatic *Chchd10*^*S59L/+*^ and wild-type mice. The mean age of mice was 43.5 ± 2.4 weeks, and percentage of females was 33% at the symptomatic stage. Tissue samples were immediately flash frozen in liquid nitrogen and stored at - 80 °C until further analyses. Tissues were ground to powder by using liquid nitrogen cooled mortar and pestle. 11.1 mg ± 1.7, and 10.1 mg ± 2.1 of powder tissue was used for metabolite and lipid extraction, respectively. Blood was taken by cardiac puncture from mice at pre-symptomatic (16 to 20 weeks of ages, n=5 mice per group) and symptomatic (40 to 48 weeks of ages, n=7 mice per group) stages, and transferred to EDTA coated tubes (Sarstedt). Then, blood was centrifuged at 500 rpm for 10 min at 4°C, plasma was collected and immediately frozen in microtubes (Eppendorf) and stored at -80°C until further analyses.

### Metabolomics analysis

Frozen tissues were extracted with 1 mL cold acetonitrile/water (1:1, v/v). The suspension was vortexed, incubated at 4°C for 1 h, centrifuged at 16 000 x g for 10 mins at 4°C and then 450 µL of the supernatant was dried. For plasma metabolites extraction, 20 µL were precipitated with methanol and centrifuged at 16 000 x g for 10 mins at 4°C, and the supernatant was dried. All dried metabolites extract from tissues were reconstituted in 100 µL of either methanol/water (1: 9, v/v) or ACN/water (9: 1, v/v) depending on the LC-column used and 10 µL was injected into LC-HRMS system.

LC-HRMS analysis was performed using an UPLC Ultimate WPS-3000 system (Dionex, Germany) coupled to a Q-Exactive mass spectrometer (Thermo Fisher Scientific, Bremen, Germany) and operated in positive (ESI+) and negative (ESI-) electrospray ionization modes (analysis for each ionization mode) was used for this analysis. Liquid chromatography was performed using a Phenomenex Kinetex 1.7 µm XB – C18 column (150 mm x 2.10 mm) maintained at 55°C. Two mobile phase gradients were used. The gradient was maintained at a flow rate of 0.4 mL/min over a runtime of 20 min. Two different columns were used to increase the metabolic coverage. Accordingly, a hydrophilic interaction liquid chromatography (HILIC) column (150 mm x 2.10 mm, 100 Å) was also used. During the full-scan acquisition, which ranged from 58 to 870 m/z, the instrument operated at 70,000 resolution (m/z = 200). As required for all biological analyses, the pre-analytical and analytical steps of the experiment were validated by findings of Quality Control (QC) samples (mix of all the samples analyzed). Coefficients of variation [CV% = (the standard deviation/ mean) x 100], were calculated from all metabolites data and metabolites having a CV in QCs >30% were excluded from the final dataset. QCs were analyzed at the beginning of the run, every 13 samples and at the end of the run.

A targeted analysis was applied on the samples, based on a library of standard compounds (Mass Spectroscopy Metabolite Library (MSML®) of standards, IROA Technologies(tm)). The following criteria were followed to identify the metabolites: (1) retention time of the detected metabolite within ± 20 s of the standard reference, (2) exact measured of molecular mass of the metabolite within a range of 10 ppm around the known molecular mass of the reference compound, and (3) correspondence between isotopic ratios of the metabolite and the standard reference. The signal value was calculated using Xcalibur® software (Thermo Fisher Scientific, San Jose, CA) by integrating the chromatographic peak area corresponding to the selected metabolite.

### Lipidomics analysis

Lipids were extracted according to the Floch method [14]. 525 µL ice-cold methanol: water (3:1, v/v) was added directly to frozen powder tissue, and vortexed. Then, 400 µL chloroform and 275 µL water were added. The suspension was vortexed, and incubated at 4°C for 30 minutes. Then, the Mixture was centrifuged at 16 000 x g for 10 mins at 4°C. 350 µL of the lower phase (organic) layer was collected in the new tube and dried. Lipid extracts were reconstituted in 100 µL of chloroform and transferred to an LC vial with glass insert for analysis.

Reverse phase liquid chromatography was selected for separation with an UPLC system (Ultimate 3000, ThermoFisher). For cardiolipins analysis, lipid extracts were separated on a Cortex C8 150×2.1, 2.7µm column (Waters) operated at 400 µl/min flow rate (column temperature 45°C). The injection volume was 3 µL of diluted lipid extract. Eluent solutions were ACN/H2O 60/40 (V/V) containing 10mM ammonium formate and 0.1% formic acid (solvent A) and IPA/ACN/H2O 88/10/2 (V/V) containing 2mM ammonium formate and 0.02% formic acid (solvent B). The used step gradient of elution was: 0.0-5.0 min 60% B, 5.0-12.5 min 60 to 74% B, 12.5-13.0 min 70 to 99% B, 13.0-17 min 99% B, 17-17.5 min 99 to 60% B and finally the column was reconditioned at 60% B for 4 min.

For all other lipids, lipid extract were separated on a 1.7 μm C18 (150 mm□×□2.10 mm, 100 Å) UHPLC column (Kinetex, Phenomenex, Torrance, CA) heated at 55 °C. The solvent system comprised mobile phase A [isopropanol/ACN (9:1)□+□0.1% (vol/vol) formic acid□+□10 mM ammonium formate], and mobile phase B [ACN/water (6:4)□+□0.1% (vol/vol) formic acid□+□10 mM ammonium formate]; the gradient operated at a flow rate of 0.26 mL/min over a run time of 24 min. The multistep gradient was programmed as follows: 0–1.5 min-32–45% A, 1.5–5 min-45–52% A, 5–8 min-52–58% A, 8–11 min-58–66% A, 11– 14 min-66–70% A, 14–18 min-70–75% A, 18 to 21 min-75–97% A and 21–24 min-97% A. The autosampler temperature (Ultimate WPS-3000 UHPLC system, Dionex) was set at 4 °C, and the injection volume for each sample was 5 µL. The heated ESI source parameters were a spray voltage of 3.5 kV, a capillary temperature of 350 °C, a heater temperature of 250 °C, a sheath gas flow of 35 arbitrary units (AU), an auxiliary gas flow of 10 AU, a spare gas flow of 1 AU and a tube lens voltage of 60 V for C18. The instrumental stability was evaluated by multiple injections (n□=□9) of a QC sample obtained from a pool of 10 μL of all samples analyzed. This QC sample was injected once at the beginning of the analysis, then after every 10 sample injections and at the end of the run.

The UPLC system was coupled with a Q-exactive orbitrap Mass Spectrometer (thermofisher, CA); equipped with a heated electrospray ionization (HESI) probe. This spectrometer was controlled by the Xcalibur software and was operated in electrospray negative mode. MS spectra were acquired at a resolution of 70 000 (200 m/z) in a mass range of 1000-1600 m/z. 15 most intense precursor ions were selected and isolated with a window of 1 m/z and fragmented by HCD (Higher energy C-Trap Dissociation) with normalized collision energy (NCE) of 20, 30 and 40 eV. MS/MS spectra were acquired in the ion trap and the resolution was set at 35 000 at 200 m/z.

Data were reprocessed using Lipid Search 4.1.16 (ThermoFisher). In this study, the product search mode was used and the identification was based on the accurate mass of precursor ions and MS2 spectral pattern. Mass tolerance for precursor and fragments was set to 5 ppm and 8 ppm respectively. m-score threshold was selected at 5 and ID quality filter was fixed at grades A, B and C. [M-H]-adducts were searched.

### Proteomics analysis

Cardiac tissue was collected from pre-symptomatic (4 KI mice and 5 wild-type) and symptomatic mice (n=6 mice per group). Proteins were extracted from tissues powder and the concentration was determined using the Pierce BCA assay kit (Thermo Fisher Scientific). 5 µg of total protein extracts were stacked on acrylamide-SDS gels.

#### In-gel digestion

Protein spots were manually excised from the gel and distained by adding 100 mL of H2O/ACN (1/ 1). After 10 min incubation with vortexing the liquid was discarded. This procedure was repeated two times. Gel pieces were then rinsed (15 min) with acetonitrile and dried under vacuum. Each excised spot was reduced with 50 mL of 10 mM dithiothreitol and incubated for 30 min at 56°C. Alkylation was performed with 15 mL of 55 mM iodoacetamide for 15 min at room temperature in the dark. Gel pieces were washed by adding successively (i) 100 mL of H2O/ACN (1/1), repeated two times and (ii) 100 mL of acetonitrile. Next, gel pieces were reswelled in 60 mL of 50 mM NH4HCO3 buffer containing 10 ng/mL of trypsin (modified porcine trypsin sequence grade, Promega) incubated for one hour at 4°C. Then the solution was removed and replaced by 60 mL of 50 mM NH4HCO3 buffer (without trypsin) and incubated overnight at 37°C. Tryptic peptides were isolated by extraction with (i) 60 mL of 1% AF (acid formic) in water (10 min at RT) and (ii) 60 mL acetonitrile (10 min at RT). Peptide extracts were pooled, concentrated under vacuum and solubilized in 15 µL of aqueous 0.1% formic acid and then injected.

#### NanoHPLC-HRMS analysis

NanoHPLC-HRMS analysis was performed using a nanoRSLC system (ultimate 3000, Thermo Fisher Scientific) coupled to an Easy Exploris 480 (Thermo Fisher Scientific). Peptide separation was carried out using the Easy-nLC ultra high-performance LC system. 5 µl of peptides solution was injected and concentrated on a µ-Precolumn Cartridge Acclaim PepMap 100 C18 (i.d. 5 mm, 5 mm, 100 A°, Thermo Fisher Scientific) at a flow rate of 10 mL/min and using solvent containing H2O/ACN/TFA 98%/2%/0.1%. Next peptide separation was performed on a 75 mm i.d. x 500 mm (2µm, 100 A°) PepMap RSLC C18 column (Thermo Fisher Scientific) at a flow rate of 300 nL/min. Solvent systems were: (A) 100% water, 0.1%FA, (B) 100% acetonitrile, 0.1% FA. The following gradient was used t = 0 min 2% B; t = 3 min 2%B; t = 103 min, 20% B; t = 123 min, 32% B; t = 125 min 90% B; t = 130 min 90% B; (temperature was regulated at 40°C). MS spectra were acquired at a resolution of 120 000 (200 m/z) in a mass range of 375–1500 m/z with an AGC target 3e6 value of and a maximum injection time of 25 ms. The 20 most intense precursor ions were selected and isolated with a window of 2 m/z and fragmented by HCD (Higher energy C-Trap Dissociation) with a normalized collision energy (NCE) of 30%. MS/MS spectra were acquired in the ion trap with an AGC target 5e5 value, the resolution was set at 15,000 at 200 m/z combined with an injection time of 22 ms.

#### MS data reprocessing

MS data were subjected to LFQ analysis using MaxQuant v1.6.17.0 (http://www.macquant.org/) using the MaxQuant platform (v1.6.7.0) [20]. Database search of the MS/MS data was performed in MaxQuant using the Andromeda search engine against Mus Musculus reviewed UniProtKB database (June 2020) and MaxQuant contaminants database. Digestion mode was set to Trypsin/ P specificity, with a fixed carbamidomethyl modification of cysteine, and variable modifications of protein N-terminal acetylation and methionine oxidation. Mass deviation was set to 20 ppm for the first and 6 ppm for the main search and the maximum number of missed cleavages was set to 2. Peptide and site false discovery rate (FDR) were set to 0.01. The search for co-fragmented peptides in the MS/MS spectra was enabled (“second peptides” option). Proteins identification was performed using a minimum of 2 unique and razor peptides. Quantification was achieved using the LFQ (Label-Free Quantification) algorithm. Razor and unique peptides were used for LFQ quantification and the “label min. ratio count” was fixed at 2. The match between runs option was enabled, allowing a time window of 0.7 min to search for already identified peptides in all obtained chromatograms.

Statistical evaluation of the data was performed with the freely available Perseus statistics software (version 1.6.15.0). Common contaminants, reverse decoy matches and only identified by site were removed from the protein identification list. At least 2 unique peptides and razor per protein were required for a protein identification. Only proteins that were identified and quantifiable in at least five biological replicates in each group were used for relative quantification. For statistical evaluation the two-sample tests were used with Welch’s test and the FDR calculation was based on Permutation based FDR with the default settings of the Perseus statistics software.

### Metabolic pathway analysis

Metabolic pathway enrichment analysis and pathway topology analysis were used to identify *CHCHD10*^*S59L*^ mutation-dependent changes in global biochemical domains. MetaboAnalyst computational platform was used to test for significant enrichment in KEGG pathways (http://www.genome.jp/kegg/) among the noted metabolic perturbations and computes a single *P value* for each metabolic pathway. Significant enrichment was assessed based on the false discovery rate-adjusted hypergeometric test statistic (*p* ≤ 0.05). Pathway topology analysis applies graph theory to measure a given experimentally identified metabolite’s importance in a pre-defined metabolic pathway. Measurements were computed using centrality to estimate the relative importance of individual nodes to the overall network.

### Induced pluripotent stem cells (IPS) differentiated into human cardiomyocytes

Human iPSC clone used in this paper was the same as used in our previous study [7]. It was generated from human skin fibroblasts (69-year-old woman) and characterized as previously described [7]. To produce human cardiomyocytes from pluripotent stem cells (hiPSC-CMs), human iPSCs were differentiated into hiPSC-CMs using a small-molecule cardiac differentiation protocol and characterized as previously described [15]. Briefly, iPSCs were plated on 6-well plates coated with geltrex (Life Technology) and cultured in TeSR-E8 medium (Stemcell Technology). For hiPSC-CMs differentiation, we used RPMI 1960 medium (Life Technology) containing ascorbic acid (Sigma Aldrich) and B-27 Minus Insulin (Life Technology) (CDM). At 80% iPSC confluency, on Day 0 (D0) differentiation was initiated with the addition of the CDM medium supplemented with 6 μM CHIR-99021 (Selleckchem). After 24h of exposure to CHIR-99021, medium was changed to CDM (D1). On D3 medium was changed to 1:1 mix of spent and fresh CDM with 5 μM IWP-2 (Selleckchem). On D5, after 48h of exposure to IWP-2, the medium was change to CDM. On D7, medium was changed to RPMI 1960 with ascorbic acid and B-27 Supplement 50X (CCM). Afterwards, CCM was changed every two days until further analysis. Contracting cells were noted from d7.

For immunofluorescence analysis, on day 23, hiPSC-CMs were dissociated with TrypLE Express (Life Technologies) and reseeded on geltrex-coated plates for 5 days. After 72 treated with Acetyl-L-carnitine, cells were loaded with 100nM Mitotracker Red (MitoRed, 37°C, 10 min) to label mitochondria. Image stacks were acquired with and observed on a laser scanning confocal microscope Nikon A1R. Confocal images were acquired with a Apochromat 40x/1,25 NA oil objective lens and were collected using 561 nm laser lines for excitation and appropriate emission filters be performed on a confocal microscope (Nikon A1R) with a x40 oil immersion lens.

For electronical microscopy cells were fixed with 1.6% glutaraldehyde in 0.1 M cacodylate buffer (pH 7.4) for 2 hours, rinsed and postfixed in 1% osmium tetroxide and 1% potassium ferrocyanide in 0.1M cacodylate buffer before to processed for ultrastructure as previously described [7] and analyzed using a JEOL JEM1400 transmission electron microscope (JEOL, Tokyo, Japan).

### RNA extraction and gene expression assays

Total RNA was isolated from 10-50 mg tissues using TRIzol^®^ reagent (Invitrogen Life Technologies, USA) following the manufacturer’s instructions. 2ug of total RNA was transcribed into cDNA using Superscript synthesis system for RT-qPCR (Invitrogen) following the manufacturer’s directions.

Finally, real-time PCR was performed using the StepOnePlus Real-Time PCR system (Applied Biosystems) with SYBR Green (Applied Biosystems). The primers pairs used are listed in supplementary Table 1, the housekeeping gene RPLP0 was used as a control for RNA quality, and used for normalization. A melting curve was subsequently recorded to confirm the identity of amplified products. We employed the ΔΔCt method [16] to determine relative differences in transcript levels. We defined the mean transcript levels for wildtype mice as 1 and transcript levels of KI mice were plotted as “Fold change in transcript level” with respect to the transcript level in control mice (=1). Data are expressed as mean ± SEM. Differences in the calculated means between groups were assessed using the student’s t test.

### Western Blotting

Tissue powders or cells were lysed in lysis buffer (SDS 20%, TRIS 0.5 M pH6.8, Glycerol 100%) and the concentration of proteins was determined using the Pierce BCA assay kit (Thermo Fisher Scientific). 30 µg of total protein extracts were separated on acrylamide-SDS gels and transferred to PVDF membranes (Millipore). After the nonspecific binding sites had been blocked, the membrane was incubated overnight at 4°C with the primary antibody (listed in Supplementary Table 2). The membrane was washed 3 times and incubated with the HRP-conjugated secondary antibody for 1 hour at room temperature. Immunoblots were visualized using the enhanced chemiluminescence detection kit (Pierce).

### Statistical analysis

Univariate and multivariate analyses were performed using metaboanlyst platform. Univariate analysis was performed by *t*-test and fold change analysis. Multivariate analysis was performed by principal component analysis (PCA) to validate the quality of analytical system and to observe possible outliers. Meanwhile, Orthogonal partial least squares-discriminate analysis (OPLS-DA) was applied to establish a relational model between experimental samples and the higher abundance amount to create a model prediction for different samples, and the results were visualized in score plots to display the group clusters. The Q2 (predicted variation) and R2 (explained variation) parameters were used to evaluate the models. In addition, the corresponding influence intensity and explanation capacity of each metabolite’s higher abundance mode effects for sample groups was estimated by the value of variable importance of projection (VIP).

### Resource availability

Metabolomics data from heart tissues and plasma were deposited to the Metabolomics Workbench [17] with the study ID: ST002434 and ST002435, respectively.

## Results

### *CHCHD10*^*S59L*^ mutation is associated with metabolic changes in plasma

#### Biochemistry characterization of Chchd10^S59L/+^ mice

The phenotypical characteristics and selected biochemical indexes of mice measured on plasma and collected between 25 and 33 weeks of age are presented in Table 1. Male *Chchd10*^*S59L/+*^ mice had significantly lower body fat than control littermates (4.3g ± 0.4 *vs* 9.8g ± 0.9; *p* < 0.0001). The levels of urea [(6.3g ± 0.2 *vs* 7.7g ± 0.9; *p* < 0.01) for male and (6.3g ± 0.5 *vs* 8.3g ± 0.3; *p* < 0.01) for female], creatinine [(5.0g ± 0.3 *vs* 8.1g ± 0.5; *p* < 0.0001) for male and (8.1g ± 0.4 *vs* 10g ± 0.5; *p* < 0.01) for female], lactate [(4.9g ± 0.2 *vs* 6.1g ± 0.5; *p* < 0.05) for male and (4.5g ± 0.1 *vs* 5.2g ± 0.3; *p* < 0.05) for female] and some electrolytes such as sodium (Na), potassium (K), chlorine (Cl), were significantly higher while those of leptin [(6.1g ± 1 *vs* 1.3g ± 0.4; *p* < 0.0001) for male and (3.4g ± 0.6 *vs* 0.9g ± 0.2; *p* < 0.001) for female] was significantly lower in *Chchd10*^*S59L/+*^ mice compared to wild type. Male KI mice had significantly lower total and HDL cholesterol levels than controls (1.7 mmol/L± 0.1 *vs* 2.2 mmol/L ± 0.1; *p* < 0.01), whereas their triglyceride levels were significantly higher (1 mmol/L ± 0.1 *vs* 0.7 mmol/L ± 0.04; *p* < 0.01) (Table 1). Overall indicating the *CHCHD10*^*S59L*^ mutation is associated with a global metabolic changes in the mice.

**Table 1:**
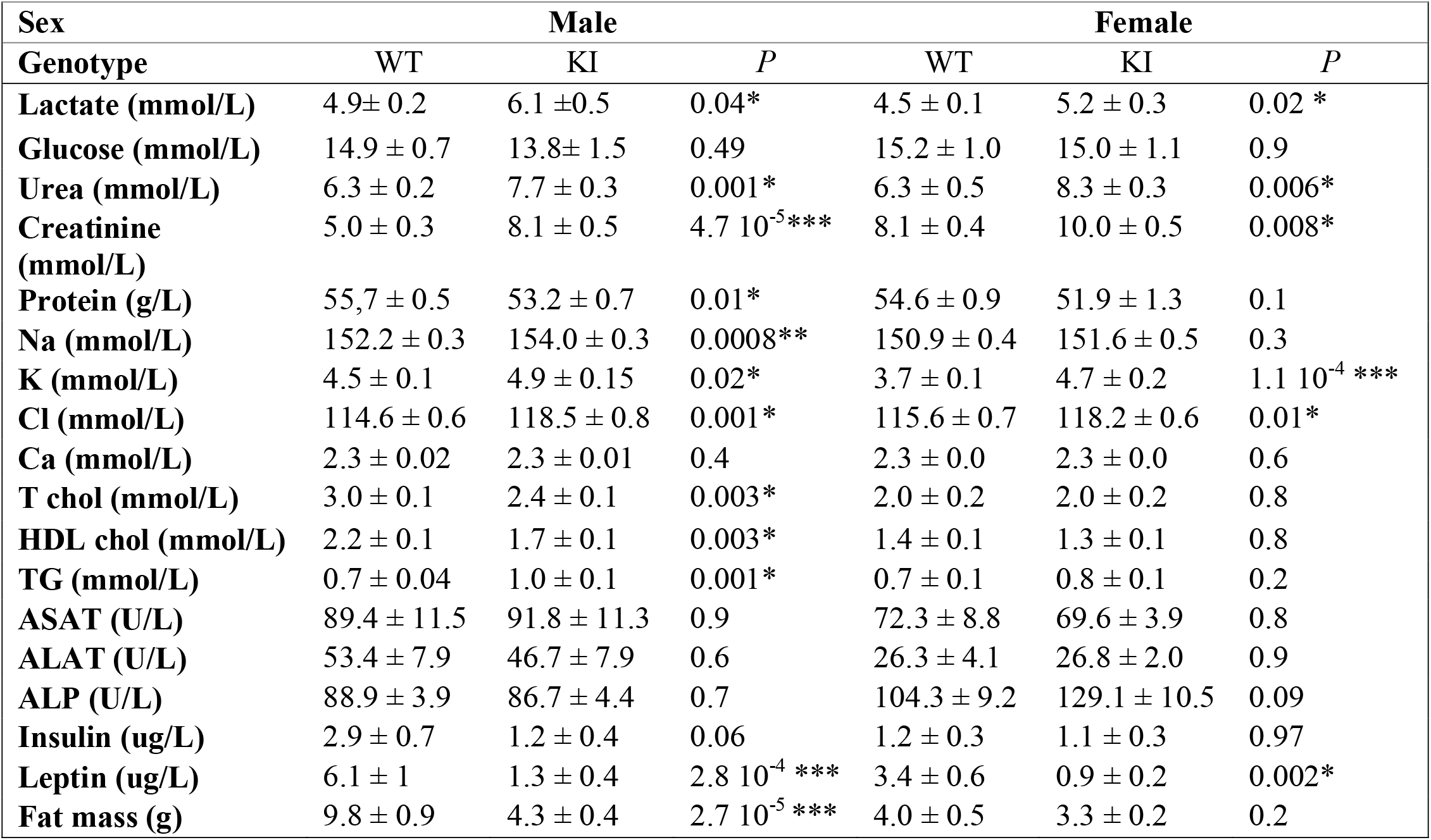
Baseline characteristics of CHCHD10^S59L^ mice and their WT littermate at 25 weeks for qNMR measurement (fat mass) and 33 weeks for biochemistry measurement (n = 9-10 mice per group). Data are expressed as mean ± SEM, *P* value < 0.05 (*); < 0.001 (**); < 0.0001(***) .Na: sodium; K: potassium; Ca: calcium; Cl: Chloride; TG: Triglyceride; ASAT: aspartate aminotransferase; ALAT: alanine aminotransferase; ALP: alkaline phosphatase; T chol: total cholesterol; HDL chol: high density lipoprotein cholesterol.

| Sex | Male |  |  | Female |  |  |
| --- | --- | --- | --- | --- | --- | --- |
| Genotype | WT | KI | *P* | WT | KI | *P* |
| Lactate (mmol/L) | 4.9 ± 0.2 | 6.1 ± 0.5 | 0.04* | 4.5 ± 0.1 | 5.2 ± 0.3 | 0.02 * |
| Glucose (mmol/L) | 14.9 ± 0.7 | 13.8 ± 1.5 | 0.49 | 15.2 ± 1.0 | 15.0 ± 1.1 | 0.9 |
| Urea (mmol/L) | 6.3 ± 0.2 | 7.7 ± 0.3 | 0.001* | 6.3 ± 0.5 | 8.3 ± 0.3 | 0.006* |
| Creatinine (mmol/L) | 5.0 ± 0.3 | 8.1 ± 0.5 | 4.7 10<sup>-5</sup>*** | 8.1 ± 0.4 | 10.0 ± 0.5 | 0.008* |
| Protein (g/L) | 55.7 ± 0.5 | 53.2 ± 0.7 | 0.01* | 54.6 ± 0.9 | 51.9 ± 1.3 | 0.1 |
| Na (mmol/L) | 152.2 ± 0.3 | 154.0 ± 0.3 | 0.0008** | 150.9 ± 0.4 | 151.6 ± 0.5 | 0.3 |
| K (mmol/L) | 4.5 ± 0.1 | 4.9 ± 0.15 | 0.02* | 3.7 ± 0.1 | 4.7 ± 0.2 | 1.1 10<sup>-4</sup> *** |
| Cl (mmol/L) | 114.6 ± 0.6 | 118.5 ± 0.8 | 0.001* | 115.6 ± 0.7 | 118.2 ± 0.6 | 0.01* |
| Ca (mmol/L) | 2.3 ± 0.02 | 2.3 ± 0.01 | 0.4 | 2.3 ± 0.0 | 2.3 ± 0.0 | 0.6 |
| T chol (mmol/L) | 3.0 ± 0.1 | 2.4 ± 0.1 | 0.003* | 2.0 ± 0.2 | 2.0 ± 0.2 | 0.8 |
| HDL chol (mmol/L) | 2.2 ± 0.1 | 1.7 ± 0.1 | 0.003* | 1.4 ± 0.1 | 1.3 ± 0.1 | 0.8 |
| TG (mmol/L) | 0.7 ± 0.04 | 1.0 ± 0.1 | 0.001* | 0.7 ± 0.1 | 0.8 ± 0.1 | 0.2 |
| ASAT (U/L) | 89.4 ± 11.5 | 91.8 ± 11.3 | 0.9 | 72.3 ± 8.8 | 69.6 ± 3.9 | 0.8 |
| ALAT (U/L) | 53.4 ± 7.9 | 46.7 ± 7.9 | 0.6 | 26.3 ± 4.1 | 26.8 ± 2.0 | 0.9 |
| ALP (U/L) | 88.9 ± 3.9 | 86.7 ± 4.4 | 0.7 | 104.3 ± 9.2 | 129.1 ± 10.5 | 0.09 |
| Insulin (ug/L) | 2.9 ± 0.7 | 1.2 ± 0.4 | 0.06 | 1.2 ± 0.3 | 1.1 ± 0.3 | 0.97 |
| Leptin (ug/L) | 6.1 ± 1 | 1.3 ± 0.4 | 2.8 10<sup>-4</sup> *** | 3.4 ± 0.6 | 0.9 ± 0.2 | 0.002* |
| Fat mass (g) | 9.8 ± 0.9 | 4.3 ± 0.4 | 2.7 10<sup>-5</sup> *** | 4.0 ± 0.5 | 3.3 ± 0.2 | 0.2 |

#### Plasma metabolic profile differences associated to CHCHD10^S59L^ mutation

In order to obtain a more precise characterization of the metabolic status of mice, we performed metabolomics analysis of plasma from *Chchd10*^*S59L/+*^ mice and their wild-type littermates at pre-symptomatic (16-20 weeks) and symptomatic (40 to 46 weeks of ages) stages. The LC-HRMS analysis provided 196 metabolites in plasma (Table 2). The plasma metabolite profiles distinguished the *Chchd10*^*S59L/+*^ mice from wild type controls using an orthogonal partial least squares-discriminate analysis (OPLS-DA) model in both the pre-symptomatic and symptomatic stages (R^2^Y= 98.9 %, Q^2^= 38.9%) (**Fig. 1A**). Eighty-eight metabolites were identified as the key metabolites [variable importance of projection (VIP) > 1] to discriminate the *Chchd10*^*S59L/+*^ mice from wild type controls at the pre-symptomatic and symptomatic stage (**Supplementary Table. S3**). Using univariate analysis (Wilcoxon test), 11 metabolites (5.6%) were significantly (*p* < 0.05) regulated in *Chchd10*^*S59L/+*^ mice compared to wild-type at pre-symptomatic and 49 metabolites (25%) significantly (*p* < 0.05) altered at symptomatic stage (**Fig. 1B, Supplementary Table. S4 and S5**). Pathway enrichment analysis showed that metabolites were enriched in amino acid metabolism, oxidation of fatty acid and carnitine biosynthesis (**Fig. 1C**). Very interestingly, most of the amino acids detected in the plasma were increased in *Chchd10*^*S59L/+*^ mice at both stages (**Fig. 1D, Supplementary Table. S4-5**), as well as carnitine, O-acetyl-L-carnitine and deoxycarnitine increased at symptomatic stage (**Fig. 1F**). Whereas cholesteryl acetate, several glycerophosphatidylcholines, and fatty acyls were decreased in *Chchd10*^*S59L/+*^ mice at the symptomatic stage (**Fig. 1E, Supplementary Table. S5**).

**Table 2:** Summary of metabolomics, lipidomics and proteomics analysis of the plasma and heart. Total number of metabolites, lipids and proteins identified in each sample type, and the number in brackets refers to number of metabolites, lipids or proteins significantly different (*P* < 0.05; T-test) between *Chchd10*^*S59L/+*^ mice and wild-type at pre-symptomatic and symptomatic stage.

|  | Stage of disease | Metabolites<br>(*P*<0.05) | Lipids<br>(*P*<0.05) | Proteins<br>(*P*<0.05) |
| --- | --- | --- | --- | --- |
| **Plasma** | Pre-symptomatic | 196 (11) |  |  |
|  | Symptomatic | 196 (49) |  |  |
| **Heart** | Pre-symptomatic |  |  | 943 (352) |
|  | Symptomatic | 156 (98) | 287 (173) | 1037 (675) |

**Figure 1:**
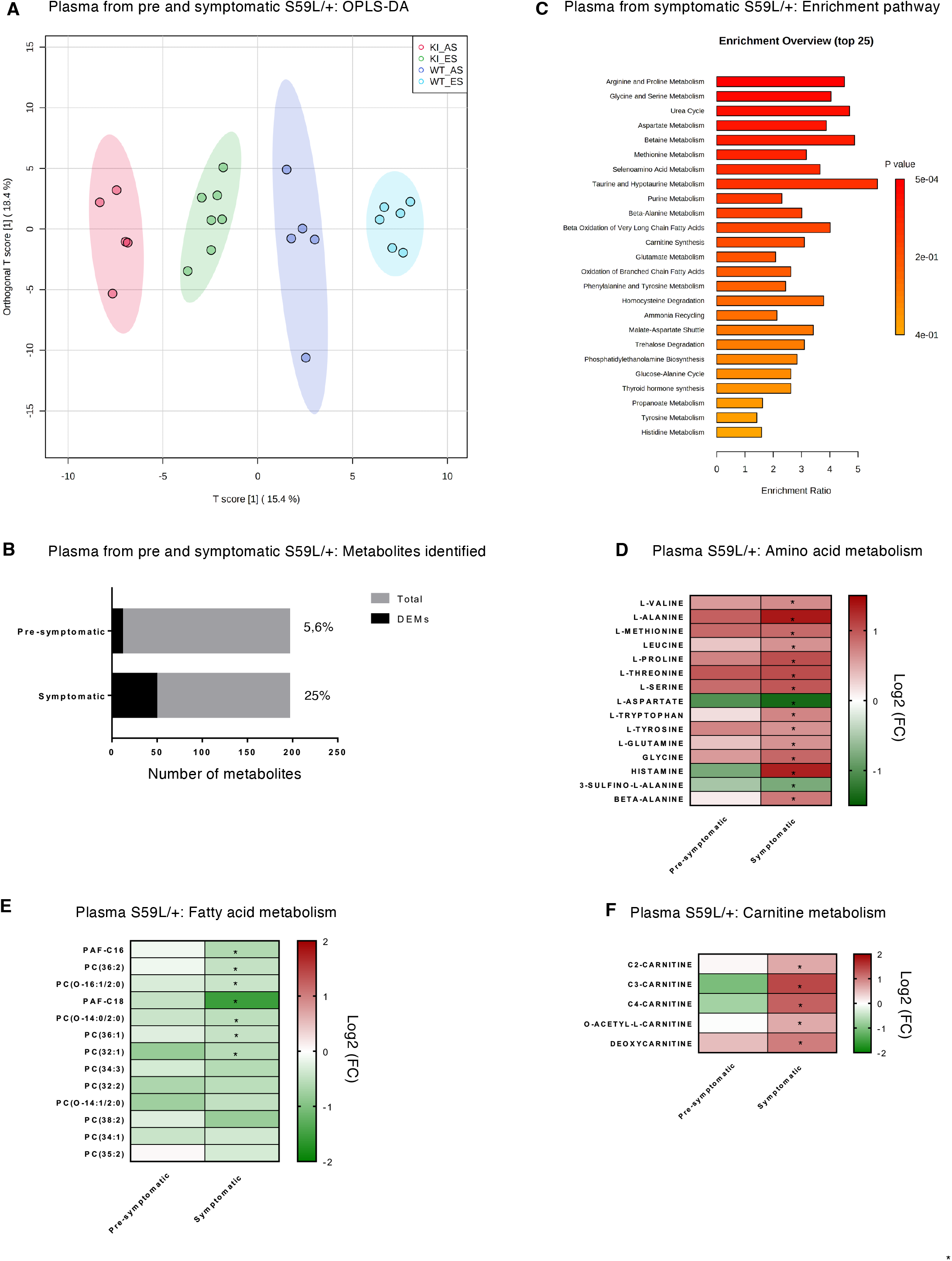
Metabolomics analysis of *Chchd10*^*S59L/+*^ plasma reveals clear group segregation and energetic metabolism alteration. (A) Score plot of the orthogonal partial least squares analysis (OPLS-DA) of plasma collected from wild-type (WT) or *Chchd10*^*S59L/+*^ (HT) mice at the pre-symptomatic (AS) and symptomatic stages (ES), (B) Number of total identified metabolites and differentially expressed metabolites (DEMs) in plasma of pre-symptomatic and symptomatic KI mice. (C) Pathway enrichment analysis was performed based on metabolite changes in plasma at the symptomatic stage. Bar colors reflect the significance (*p* value) for the specific pathway. The length of each bar indicates the impact value of each metabolic pathway. Heatmaps of the expression of plasma metabolites [represented as log_2_ (fold change)] associated with (D) amino acid, (E) carnitine and (F) fatty acid metabolism pathways, comparing *Chchd10*^*S59L/+*^ *versus* wild-type pre-symptomatic and symptomatic mice. Statistical significance was evaluated by t-test (\**p* < 0.05) and the fold change was set to >1 (more detailed in Table S2).

### Proteomics analysis reveals OXPHOS downregulation in *Chchd1010*^*S59L/+*^ heart associated with mitochondrial cardiomyopathy markers

To assess the influence of the *CHCHD10*^*S59L*^ mutation at the level of global protein expression, longitudinal proteomics were conducted to profile the expression of several proteins in *Chchd10*^*S59L/+*^ heart at pre-symptomatic and symptomatic stages. We choose to analyze the proteome in the heart as cardiomyopathy is a characteristic of *Chchd10*^*S59L/+*^ mice [7]. We performed a principal component analysis (PCA) on the differential protein expressed between the hearts of *Chchd10*^*S59L/+*^ mice and their wild-type littermate at both stages. PCA showed clear discrimination of samples by genotype and disease stage (**Fig. 2A**). Several differentially expressed proteins are observed in the hearts of mice at both stages. We observed that 208 (22%) and 483 (47%) proteins were significantly (with adjusted p < 0.05) deregulated in the heart of *Chchd10*^*S59L/+*^ mice at the pre-symptomatic and symptomatic stages, respectively (**Fig. 2B**). The 170 significantly altered proteins in heart at the pre-symptomatic and symptomatic stages (**Fig. 2C**) were subjected to gene ontology annotations of biological processes and revealed a significant enrichment of biological processes associated with mitochondrial metabolism: “oxidative phosphorylation,” “cellular respiration-ATP metabolic process,” “mitochondrial membrane,” “TCA cycle,” “lipid oxidation,” and “mitochondrial abnormality”.

**Figure 2:**
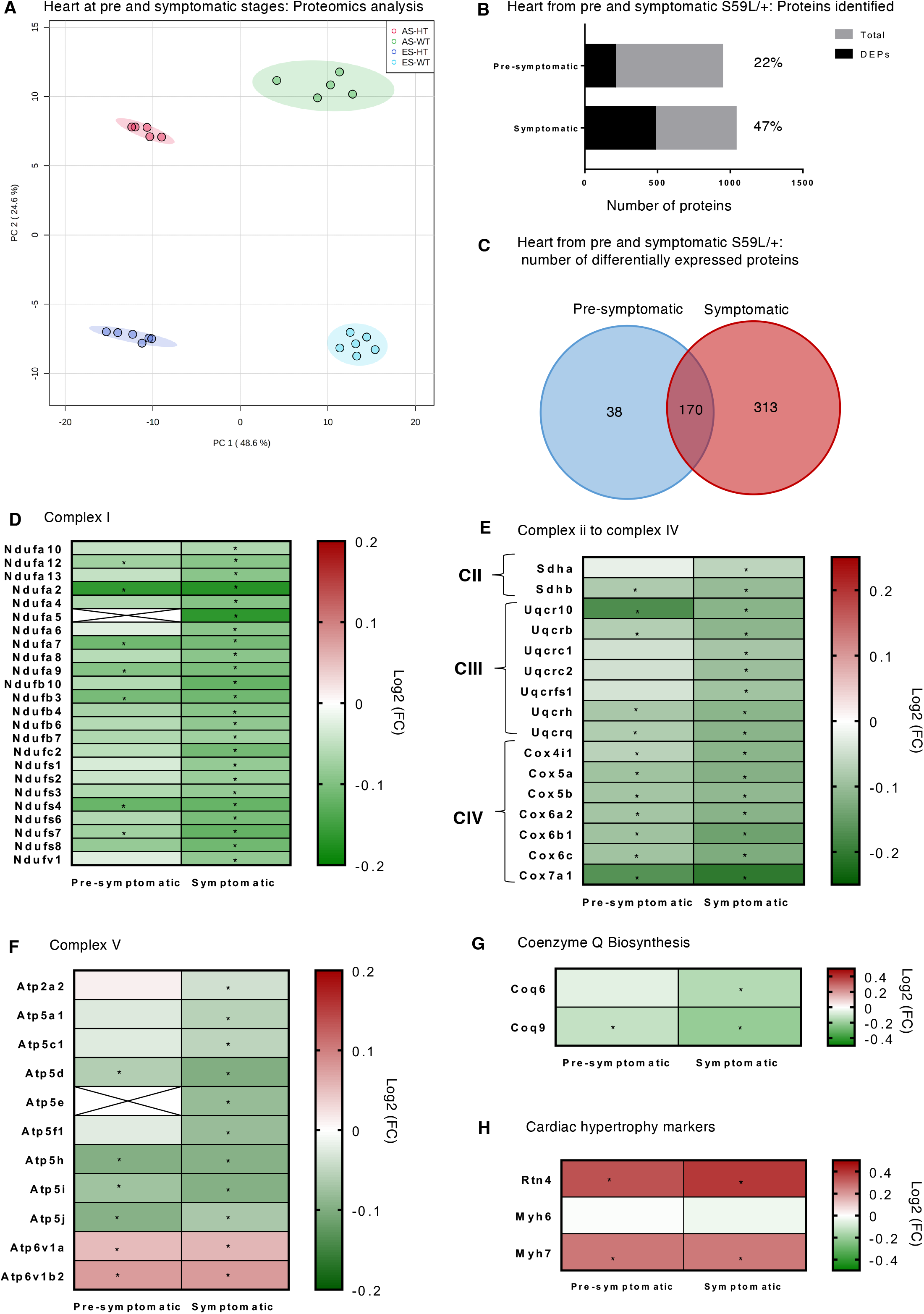
*Chchd10*^*S59L/+*^ heart presents cardiomyopathy and mitochondrial oxidative metabolism downregulation. (A) Principal component analysis (PCA) showing differential protein expression profiles between wild-type (WT) and *Chchd10*^*S59L/+*^ heart (HT) at pre-symptomatic (AS) and symptomatic stages (ES). (B) Number of total identified proteins and differentially expressed proteins (DEPs) at both stages. (C) Venn diagram of proteins significantly (with adjusted p value < 0.05) altered in hearts of *Chchd10*^*S59L/+*^ mice at pre-symptomatic and symptomatic stages. Heatmaps of proteins expression [represented as log_2_ (fold change)] associated with (D-F) OXPHOS complexes I-V, (G) CoQ biosynthesis and (H) cardiomyopathy markers, comparing *Chchd10*^*S59L/+*^ *versus* wild-type at the pre-symptomatic and symptomatic stages. Statistical significance was evaluated by t-test (* with adjusted p < 0.05).

Consistent with the oxidative phosphorylation defects previously described [7], we observed a decrease in most respiratory chain complex I-IV and complex V subunits at both stages (**Fig. 2D-F)**. We find that the lipid-binding proteins (Coq9 and Coq6), involved in the biosynthesis of coenzyme Q (an essential lipid-soluble electron transporter), were also significantly downregulated in the heart of symptomatic *Chchd10*^*S59L/+*^ mice compared to their wild-type littermates (**Fig. 2G)**.

Accordingly to the mitochondrial cardiomyopathy phenotype of *Chchd10*^*S59L/+*^ mice previously described [7], we observed an increase in protein levels of hypertrophic cardiomyopathy markers such as myosin heavy-chain beta (MYH7) and reticulon 4 (RTN4) starting at the asymptomatic stage (**Fig. 2H)**. These data demonstrate that mitochondrial cardiomyopathy associated with OXPHOS alterations appears in the early stage of disease progression in *Chchd10*^*S59L/+*^ mice model.

### β-oxidation impairment is associated with a strong modulation of lipid metabolism in *Chchd10*^*S59L/+*^ heart

Proteomics analysis reveals also changes in the proteins content of β-oxidation and fatty acid metabolism pathways in KI mice (pre-symptomatic or symptomatic) compared to age-matched wild-type mice. Indeed, we observed that NADH-cytochrome b5 reductase (Cyb5r1), which is involved in fatty acids desaturation and elongation, is upregulated in the heart at the pre-symptomatic and symptomatic stages, whereas 3-ketoacyl-CoA thiolase (Acaa2), which is one of the enzymes that catalyzes the last step of the mitochondrial β-oxidation pathway breaking down fatty acids into acetyl-CoA, was decreased in the same tissue (**Fig.3A**). In cells, long-chain fatty acids dependent on esterification with L-carnitine to form acetyl-carnitine for transport from the cytoplasm to the mitochondrial matrix for oxidation and energy production. Accordingly, we observed that most of the transporters and enzymes involved in the β-oxidation pathway were down-regulated with few exceptions, such as an increase in phospholipase A-2-activating protein (Plaa) expression at both stages (**Fig.3A)**. To determine whether the mRNA levels of some of those transporters and enzymes were affected, we performed RT-qPCR analysis on *Chchd10*^*S59L/+*^ heart at both stages (**Fig. 3B, supplementary Fig-S1-A)**. In lipid biosynthesis pathway, protein levels of long-chain-fatty-acid-CoA ligase 1 (Acsl1; as well as mRNA expression) and mitochondrial short-chain specific acyl-CoA dehydrogenase (Acads) in the hearts of symptomatic *Chchd10*^*S59L/+*^ mice were significantly lower than those of age-matched wild-type mice (with adjusted p < 0.05) (**Fig. 3A-B**). Regarding lipid transport in mitochondria, protein levels of fatty acid binding protein 3 (Fabp3; as well as mRNA expression), phospholipase A-2-activating protein (Plaa) in the *Chchd10*^*S59L/+*^ heart were significantly lower than those of wild-type mice at the symptomatic (with adjusted p < 0.05). With respect to the lipid oxidation, protein levels of carnitine-O-Acetyltransferase (Crat; as well as mRNA expression), carnitine-palmitoyl-transferase-2 (Cpt2; as well as mRNA expression), Acetyl-CoA acetyltransferase (Acat1), Acetyl-coenzyme A synthetase (Acss1), mitochondrial carnitine/acylcarnitine carrier protein (Scl25a20) and mitochondrial 3-ketoacyl-CoA thiolase (Acaa2) were significantly lower in *Chchd10*^*S59L/+*^ than wild-type heart at symptomatic stage (with adjusted p < 0.05) (**Fig. 3B)**.

**Figure 3:**
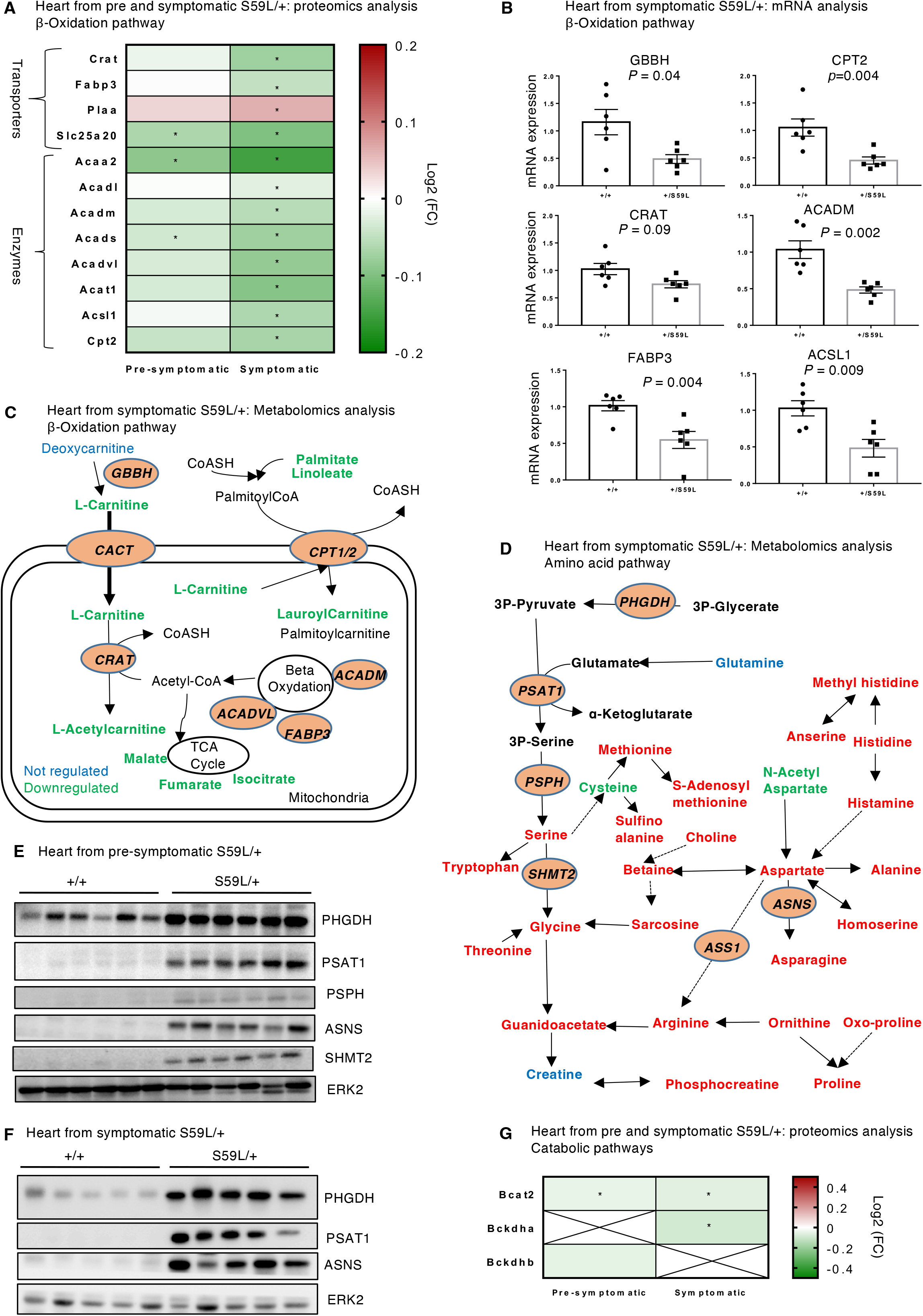
*Chchd10*^*S59L/+*^ heart exhibits β-oxidation pathway downregulation and amino acid metabolism upregulation. (A) Heatmap of proteins expression [represented as log_2_ (fold change)] associated with β-oxidation pathway in heart at pre-symptomatic and symptomatic stages. (B) Expression level of mRNAs coding for genes involved in β-oxidation pathway in heart from *Chchd10*^*S59L/+*^ and wild-type mice at symptomatic stage. Data are presented as mean ± SEM compared wild-type (arbitrarily set at 1), n=6 per group, unpaired student’s t test, relative quantification was performed calculating the ratio between the gene of interest normalized against the housekeeping gene RPLP0: ribosomal protein lateral stalk subunit P0. GBBH: gamma-butyrobetaine dioxygenase; CPTII: carnitine-palmitoyl-transferase-2; CRAT: carnitine acetyltransferase; ACADM: Acyl-CoA deshydrogenase medium chain; FABP3: fatty acid binding protein 3 and ACSL1: Acyl-coA synthethase long chain family member 1. (C) Summary of energetics metabolites identified in heart samples from mice at symptomatic stage. The substances in green were found significantly downregulated while the one in blue were not modulated. (D) Schematical pathway of amino acid metabolism pathways altered in heart of *Chchd10*^*S59L/+*^ mice at the symptomatic stage. In green (downregulated), blue (not modulated) and red (upregulated) were the metabolites identified in this pathway. Western blot showing the increased levels of some enzymes involved in the amino acid metabolism pathway in the heart of (E) pre-symptomatic (n=6 per group) and (F) symptomatic (n=5 per group) *Chchd10*^*S59L/+*^ mice. Each line represents an individual mouse. Erk2 is used as a loading control. PHGDH: Phosphoglycerate Dehydrogenase, PSAT1: Phosphoserine Aminotransferase 1, ASNS: Asparagine Synthetase, PSPH: Phosphoserine Phosphatase, SHMT2: Serine Hydroxymethyltransferase 2. (G) Heatmap of proteins expression [represented as log_2_ (fold change)] associated with catabolic pathway in heart at pre-symptomatic and symptomatic stages.

To further determine which lipids alterations are associated with the *CHCHD10*^*S59L*^ mutation, we performed a lipidomic analysis in cardiac tissue from *Chchd10*^*S59L/+*^ mice at symptomatic stage. A total of 287 lipids were identified in the heart (**Table 2, supplementary Fig.S1-B**). PCA analysis revealed that *CHCHD10*^*S59L*^ mutation had the strongest impact on the lipid profile of heart (**Supplementary Fig. S1-C**). Univariate analysis performed by one-way ANOVA yield 173 altered lipids in in heart (**Table 2, supplementary Fig.S1-D**). This analysis indicates that the most striking changes in lipids are related to a decrease in triglycerides (TGs) and an increase in esters cholesterols (ChE), ceramides and most of the phospholipids in the heart (**Supplementary Fig. S1 D-E**). This is in agreement with the measured decreases in TGs and regulation of lipid metabolism in serum and plasma of *Chchd10*^*S59L/+*^ mice compared with the wild-type (**Table 1, Supplementary Table. S4-5**). Cardiolipin alteration is observed in the heart, with approximately 80% of identified cardiolipin altered in *Chchd10*^*S59L/+*^ heart compared to wild-type (**Supplementary Fig. S1-E**). Taken together these results suggest that *Chchd10*^*S59L/+*^ mice present a disturbance of mitochondrial oxidative metabolism with lipids profile alterations in the heart and plasma.

### β-oxidation and OXPHOS proteins changes affect energy homeostasis and amino-acid metabolism in *Chchd10*^*S59L/+*^ hearts

To assess whether changes in OXPHOS, β-oxidation proteins expression affected global metabolism in heart, we performed metabolomics analysis on *Chchd10*^*S59L/+*^ heart at symptomatic stage. PCA analysis on heart metabolome from *Chchd10*^*S59L/+*^ and wild-type mice showed clear clustering of samples by genotype (**Supplementary Fig. S1-F)**. We observed decrease in TCA cycle components (fumarate, isocitrate and malate) and β-oxidation intermediates metabolites (L-carnitine, L-acetylcarnitine, lauryolcarnitine and palmitate) in *Chchd10*^*S59L/+*^ mice (**Fig. 3C**). Interestingly, a significant decrease in fatty acid biosynthesis intermediates, such as malonate and ethyl-malonate, was also observed in the hearts of symptomatic *Chchd10*^*S59L/+*^ mice (**Fig. 3C**).

Regarding the amino-acid modulation in the *Chchd10*^*S59L/+*^ heart, we observed a significant increase in methionine, phenylalanine, arginine, proline, lysine, threonine, glycine, tryptophan, serine, asparagine, aspartate, alanine and histidine (**Fig. 3D**). Consistently with the up-regulation of cardiac amino acids levels, we observed an upregulation of the protein levels of some enzymes involved in amino acid metabolism, such as a phosphoglycerate dehydrogenase (PHGDH), phosphoserine aminotransferase (PSAT1) and asparagine synthetase (ASNS), both in the heart of pre-symptomatic (**Fig. 3E**) and symptomatic mice (**Fig. 3F**).

The intra-tissue homeostasis of essential amino acids (e.g., methionine, phenylalanine, lysine, valine and threonine) is largely maintained by the balance of protein synthesis and degradation, as well as amino acid catabolic activities through a production of acetyl-CoA and succinyl-CoA feeding the TCA cycle for respiration. Catabolic enzymes are mainly located in the mitochondrial matrix such as the branched-chain alpha-keto dehydrogenase complex (Bckdha, Bckdhb) and branched-chain-amino-acid aminotransferase (Bcat2). We observed that the protein level of Bcat2 is significantly downregulated in the heart of *Chchd10*^*S59L/+*^ mice at both stages, Bckdhb and Bckdha protein levels are downregulated at pre-symptomatic and symptomatic stages, respectively (**Fig. 3G**). Altogether, those results suggest that *CHCHD10*^*S59L*^ mutation induces branched-chain amino acids catabolic defects and increased non-essential amino acids synthesis which may contribute to the elevated levels of amino acids metabolites observed in plasma and heart of *Chchd10*^*S59L/+*^ mice (**Fig.1D, Fig. 3D**). In addition, β-oxidation impairment observed in *Chchd10*^*S59L/+*^ heart is also associated with carnitine metabolism downregulation (**Fig.1C)**, this finding suggests that carnitine pathway could be a potential therapeutic target for CHCHD10-related disease.

### Targeting β-oxidation pathway with L-acetylcarnitine supplementation improves mitochondrial network in IPS-derived cardiomyocytes from patient carrying *CHCHD10*^*S59L*^ mutation

To investigate the impact of these severe metabolic changes induced by CHCHD10^S59L^ mutation in the heart, we generated IPS-derived cardiomyocytes from patient carrying the CHCHD10^S59L^ mutation and from healthy control. Our protocol gave rise to cardiomyocytes which exhibit self-contractility and other distinguishing features of functional cardiomyocytes, including formation of sarcomeric organization of myosin filaments (**Fig. 4A**) and expression of cardiac specific markers such as troponin (**Fig. 4B**). We observed loss and disorganization of myosin filaments in *Chchd10*^*S59L/+*^ iPS-derived cardiomyocytes compared to wild-type as previously observed in the heart from *Chchd10*^*S59L/+*^ mice [7] (**Fig. 4A**).

**Figure 4:**
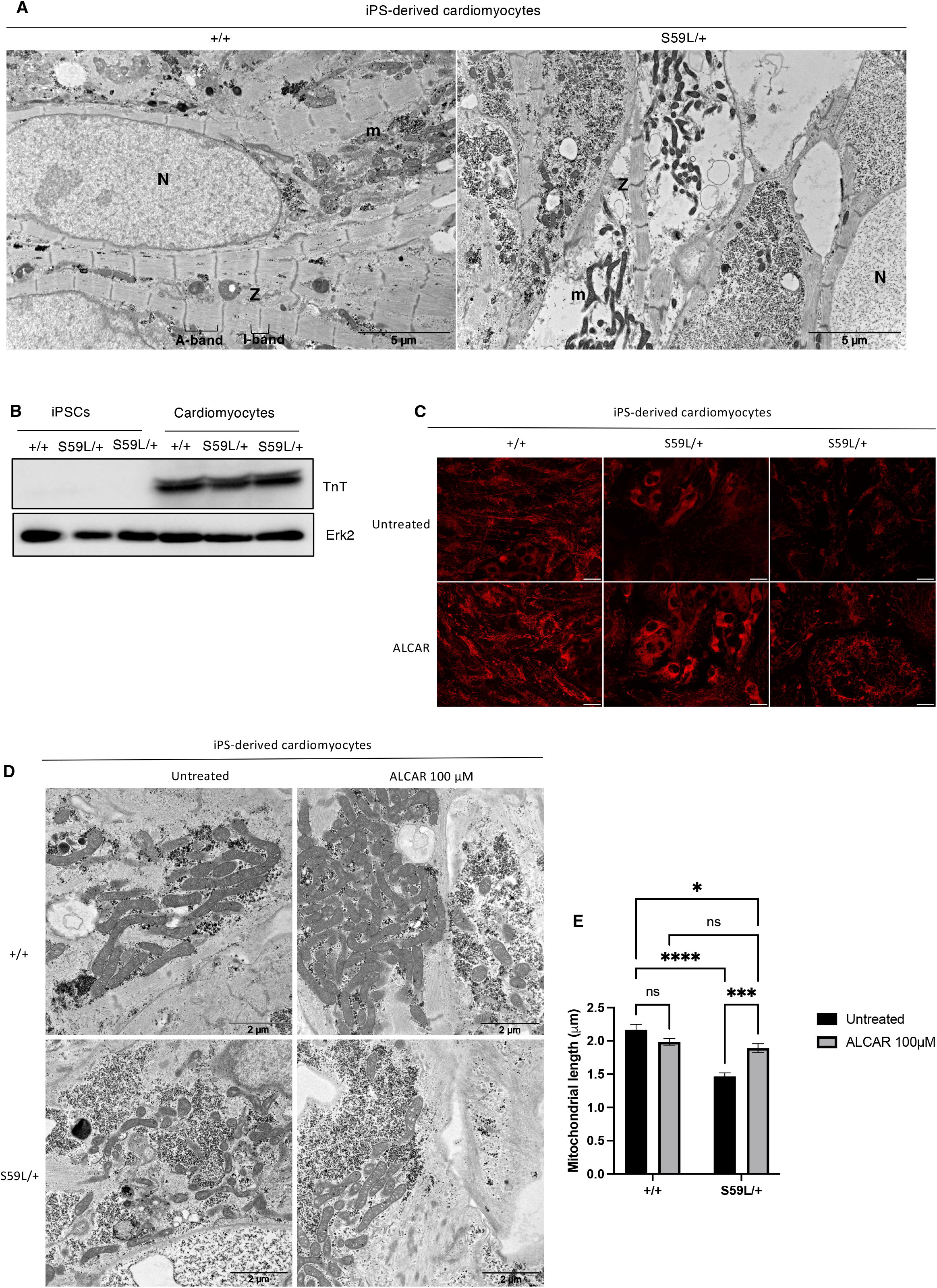
Acetyl-L-carnitine restores mitochondrial network and length in *Chchd10*^*S59L/+*^ IPS-derived cardiomyocytes. (A) iPS cell-derived cardiomyocytes from control and *Chchd10*^*S59L/+*^ patients fibroblasts were characterized by transmission electron microscopy. Ultrastructural analysis of cardiomyocytes showing the presence of initial sarcomeric organization of myosin filaments already identifiable A-and I-bands together with Z-lines. Abbreviations: *m* mitochondria, *N* Nucleus, *Z* Z-line, scale Bar 5 μm. Pictures are representative of several. (B) Western blot showing the induction of Troponin (TnT) expression in 30-day iPS cell-derived cardiomyocytes from control and *CHCHD10*^*S59L/+*^ patients. Erk2 was used as a loading control (C) Representative images of mitochondrial network from *Chchd10*^*S59L/+*^ and wild-type cardiomyocytes-derived iPSCs treated with 100 µM acetyl-L-carnitine (ALCAR) for 48h or untreated and then stained with the mitochondrial marker mitotraker Red, Scale bar: 20 μM. (D) Transmission electronic microscopy (TEM) analysis of mitochondrial shape and ultrastructure of indicated iPS cell-derived cardiomyocytes treated as in C, scale bar: 2 µM. (E) Measurement of mitochondria length of cardiomyocytes derived from S59L/+ and control iPSC (+/+) lines at day 30 of differentiation (n [100 mitochondria measured per condition) treated as in C. Two-way ANOVA, Sidak’s multiple comparisons (* p-value < 0.02; *** p-value < 0.001; ****p-value < 0.0001).

We first analyzed the mitochondrial network using the Mitotracker Red staining and the ultrastructure of mitochondria using electron microscopy. We observed that mutant cardiomyocytes exhibited a fragmented mitochondrial network (**Fig. 4C**) with shorter mitochondrial length and disorganized mitochondria cristae (**Fig. 4D-E**) compared to wild type. Next, we evaluated the potential of L-acetylcarnitine to rescue the mitochondrial phenotype associated with *CHCHD10*^*S59L*^ based on our metabolomics data showing the downregulation of carnitine metabolites in heart from *Chchd10*^*S59L/+*^ mice. For this purpose, we supplemented IPS-derived cardiomyocyte culture with acetyl-L-carnitine for 72 hours and observed a reduction in mitochondrial morphology abnormalities including a rescue of mitochondrial length and cristae phenotype in cardiomyocytes from patients with a *CHCHD10*^*S59L*^ mutation (**Fig. 4 C–D**). We showed that L-acetylcarnitine significantly restores mitochondria length from *Chchd10*^*S59L/+*^ cardiomyocytes (**Fig.4D, E**). Those data support the potential therapeutic efficacy of L-acetylcarnitine treatment on mitochondrial dysfunction induced by *CHCHD10*^*S59L*^ mutation.

## Discussion

### *Chchd10*^*S59L/+*^ mice display ALS phenotype and characteristic plasma biomarkers

In this study, we performed a comprehensive “multiple-omics” analysis of CHCHD10^S59L^ related ALS disease using *Chchd10*^*S59L/+*^ mice that we recently generated [7]. Control of whole-body energy homeostasis, i.e., the balance between energy uptake and expenditure, is crucial for maintaining stable body weight and thus overall health. We have previously observed that *Chchd10*^*S59L/+*^ mice failed to gain weight normally and were significantly leaner than controls [7]. In this study, we found also that *Chchd10*^*S59L/+*^ mice displayed a decrease body fat and lower levels of circulating leptin (**Table 1**). Leptin is produced primarily by adipose tissue and known to regulate body weight [18], its concentration in plasma is typically proportional to fat mass [19]. *In vivo* administration of leptin to normal weight has been showed to induce inhibition of food intake, suppression of lipogenesis, increase of triglyceride hydrolysis and decrease body fat stores, consistent with a role for leptin in the control of body weight [19, 20]. Increases in leptin secretion from adipocytes have been also linked to insulin stimulation [21], however we didn’t observe significantly increased insulin concentration in plasma *Chchd10*^*S59L/+*^ mice. Interestingly, we found a significantly decrease in circulating creatinine level (**Table 1**) for both male and female *Chchd10*^*S59L/+*^ mice. Plasma creatinine has been described as a prognostic biomarker for ALS, lower baseline creatinine levels was associated with a short survival in both male and female ALS patients [22-24]. Circulating contents of cholesterol, triglycerides and fatty acids were also found to be strongly altered in male *Chchd10*^*S59L/+*^ mice (**Table 1**), those alterations are known to occur over the course of ALS reflecting the metabolic environment [25]. This study showed that *CHCHD10*^*S59L*^ mutation is related to ALS phenotype with some specific known plasma biomarkers which correlated with described pathomechanisms of ALS [26, 27].

### *Chchd10*^*S59L/+*^ heart exhibits OXPHOS deficiency and cardiomyopathy markers

Mitochondria are essential cell organelles in the heart, they provide 90% of the energy required through OXPHOS and occupy 30% of cardiomyocytes volume [13]. Previously, we have showed that *Chchd10*^*S59L/+*^ mice develop a fatal mitochondrial cardiomyopathy with mtDNA instability and OXPHOS deficiency in muscle and heart [5, 7]. Consistent with OXPHOS downregulation, proteomics analysis of *Chchd10*^*S59L/+*^ hearts at pre- and symptomatic stages confirmed a dramatic decrease in protein levels of most respiratory chain complex I-V subunits (**Fig. 2D-F)**. Similar results have been found in hearts of a similar mouse model of mitochondrial cardiomyopathy [28], highlighting an early mitochondrial abnormalities in CHCHD10-related cardiomyopathy progression.

Alterations of metabolic pathways have been shown to cause a severe dysregulation of metabolites that contributes to hypertrophic cardiomyopathy and heart failure [12]. In this respect, *Chchd10*^*S59L/+*^ mice showed increased protein levels of hypertrophic cardiomyopathy markers such as myosin heavy-chain beta (MYH7) and reticulon 4 (RTN4) (**Fig. 2H**), starting in pre-symptomatic stage. The ubiquitin C-terminal hydrolase isozyme L1 (Uchl1), member of the ubiquitin proteasome system (UPS) was increased in *Chchd10*^*S59L/+*^ mice heart at both stages compared to wild-type. Numerous studies have demonstrated that UPS plays an important role in the pathogenesis of cardiac fibrosis. It has been shown to be increased significantly in infarct hearts and cardiac fibroblasts stimulated by TGF-β1 [29]. These data are corroborated by a recent study showing increased cardiac stress markers (MYH7, MYH6 and RTN4) early in disease progression [28], suggesting that *Chchd10*^*S59L/+*^ mice could represent a suitable model to investigate the pathogenesis of mitochondrial cardiomyopathies associated with OXPHOS deficiencies.

### *CHCHD10*^*S59L*^ mutation is associated with a strong alteration of the lipidome in heart

Metabolic remodeling, changes in mitochondrial structure or function, decreases the energy supply by mitochondria. Lipids play an essential role in the structuring of mitochondria. The tight association with the mitochondrial phospholipid cardiolipin (CL) ensures its structural integrity and coordinates enzymatic activity. To study lipidome alterations in the heart of *Chchd10*^*S59L/+*^ mice, we used a targeted approach by analyzing specific lipid species. Our results showed *CHCHD10*^*S59L*^ mutation is associated with modified lipid profile in heart as observed by PCA (**Supplementary Fig. S1C**). The major lipidome alterations in heart were related to cholesteryl esters, glycerophospholipids, triglyceride and cardiolipins (**Supplementary Fig. S1C-D**). We observed a decrease of circulating lipids (phospholipids) in *Chchd10*^*S59L/+*^ mice plasma (**Fig. 1E**), while some amino acids (tryptophan, methionine, tyrosine, and proline) and carnitine derivatives were significantly increased at both stages (**Fig. 1D and 1F**) indicating imbalance between the input (diet, proteolysis) and the output (protein synthesis, catabolism for energy). Elevated levels of amino acids and their intermediaries in plasma or tissues have also been associated with cardiovascular disease [30, 31].

As highlighted by our study, *Chchd10*^*S59L/+*^ mice displayed markedly elevated levels of cholesteryl esters, some glycerophospholipids and reduced concentrations of triglycerides species in heart at the symptomatic stage. Changes in cholesteryl ester metabolism were not reflected in changes in plasma total cholesterol pools (**Table 1**). In other words, it appears that cholesterol synthesis is upregulated but does not result in accumulation of free cholesterol. Abnormal mitochondrial morphology has been observed in various familial ALS models [6, 32]. Reflecting mitochondrial dysfunctional, our data reveal a significant alteration (up- and down-regulated) of cardiolipin levels in *Chchd10*^*S59L/+*^ heart at the symptomatic-stage compared with match-age wild-type group. Cardiolipin is the major phospholipid of mitochondria, specifically located in the inner membranes. The functions of cardiolipin were mainly related to ATP production, curvature stress control and mitochondria cristae morphology [33]. Given its function in structuring mitochondrial function and morphology, we suggest that decreased cardiolipin levels may partially reflect the loss and disorganization of mitochondrial cristae as previously observed [7] and thus dysfunctional mitochondria in the heart of symptomatic *Chchd10*^*S59L/+*^ mice.

Lipid alterations in ALS have been studied mainly by targeted analysis of some specific lipid species. Several studies have revealed increased levels of sphingolipids and cholesteryl esters in the spinal cord of the ALS mouse model [25]. It has been shown that triacylglyceride metabolism and glycerophospholipid and sphingolipid biosynthesis are associated with progression rate in ALS [34]. Our data support previous reports suggesting that alteration of some specific lipid species such as cardiolipin, cholesteryl esters and triglycerides could be markers of hypermetabolism in ALS. Furthermore, these results confirm that lipids metabolism could play an essential role in the relationship between hypermetabolism and CHCHD10 related disease.

### Reduced intermediate levels of carnitine metabolism in *Chchd10*^*S59L/+*^ heart are associated with impaired lipid biosynthesis and oxidation

L-Carnitine is a key molecule in both mitochondrial and peroxisomal lipid metabolism [35]. The mitochondrial carnitine system plays a crucial role in β-oxidation of long-chain fatty acids by catalyzing their transport into the mitochondrial matrix [36]. We observed increased levels of carnitine, o-acetyl-L-carnitine and deoxycarnitine in the plasma of *Chchd10*^*S59L/+*^ mice (**Fig. 1F**), whereas L-carnitine, L-acetylcarnitine and lauryolcarnitine were decreased in the heart of symptomatic mice (**Fig. 3C**). Consistent with these data, we observed in the heart, a decrease in protein and mRNA levels of the carnitine acyltransferases (CRAT) (**Fig. 3A-B**) that catalyze the exchange of coenzyme A (CoA) with L-carnitine on acyl moieties and carnitine palmitoyltransferase 2 (CPT2) (**Fig. 3A-B**), which is required for the transport of acyls into the mitochondrial matrix and their subsequent oxidation. In addition to the downregulation of the carnitine pathway in symptomatic *Chchd10*^*S59L/+*^ heart, we showed impairment of lipid biosynthesis pathways (decrease protein level of Acsl1, Gpx and Acads) and β-oxidation (decrease protein level of Crat, Cpt2, Acat1, Acss1, Scl25a20 and Acaa2), as well as decreased levels of proteins involved in lipid transport into mitochondria such as Fabp3 (**Fig.3A**). Based on these data, we can assume that *Chchd10*^*S59L/+*^ mice are deficient in the utilization of fatty acids for energy production, which is associated with decreased levels of L-carnitine and L-carnitine-related enzymes, as demonstrated in several studies [37]. It is known that a healthy heart prioritizes fatty acid utilization through β-oxidation over glycolysis and alters substrate utilization by upregulating glycolysis under stressful conditions [38, 39]. Consistent with this hypothesis, we observed increased glycolytic activity in *Chchd10*^*S59L/+*^ mice with a higher lactate level (**Table 1**) and decreased plasma glucose levels at symptomatic stage (**Supplementary Table S5**). Interestingly, we observed an increase in the protein level of the enzyme that initiates the first step of glycolysis by catalyzing phosphorylation of D-glucose to D-glucose 6-phosphate, hexokinase-1 (HK1) in the heart at both stages. These results confirm that the *CHCHD10* mutant induces metabolic remodeling from OXPHOX to glycolysis, particularly in the heart, as recently demonstrated in *Chchd10*^*S55L/+*^ knock in mice [28].

### L-acetylcarnitine improves mitochondrial ultrastructure in human cardiomyocytes carrying CHCHD10^S59L^ mutation

L-acetylcarnitine is known to play crucial role in the uptake of acetyl-CoA into the mitochondria during fatty acid oxidation, the production of acetylcholine, and the synthesis of protein and phospholipid required for membrane formation and integrity [35]. We evaluated the therapeutic efficacy of L-acetylcarnitine treatment on mitochondrial dysfunction using human cardiomyocytes. Our study shown that treating cardiomyocytes *in vitro* with L-acetylcarnitine for 72 hours restores mitochondrial network and length abnormalities induced by *CHCHD10*^*S59L*^ mutation (**Fig. 5**).

Several studies in animal model have shown that chronic treatment with acetyl-L-carnitine has a beneficial therapeutic effect on various chronic neurological diseases by increases lifespan, improves cognitive behaviour in aged animals and improves long-term memory performance [35, 40, 41]. Moreover, because of its contribution to bioenergetic processes, it plays a major role in diseases correlated with metabolic compromise, such as mitochondrial-related disorders [42]. Increased plasma concentrations of acylcarnitines (which formed from carnitine and acyl-CoAs) are suggested as a marker of metabolism disorders related to cardiovascular diseases [43, 44]. Based on our data and the literature, we suggested that targeting carnitine metabolism pathway could counterbalance the metabolic disturbances, ameliorate mitochondrial functions, and therefore delay *CHCHD10*-related disease progression.

Combining lipidomics, metabolomics and proteomics analysis, we observed that *CHCHD10*^*S59L*^ mutation induced dramatic metabolic changes in plasma and heart with many significant changes in lipids, polar metabolites and several proteins involved in OXPHOS and β-oxidation pathways. These results provide a very useful set of data and evidence of altered energy homeostasis and lipid metabolism that could be considered as essential components of the disease process associated with the *CHCHD10* mutation. Our work also suggests that sustaining the energy homeostasis with a high energetic substrate or that targeting energy metabolism pathways to improve mitochondrial function is a promising therapeutic prospect. Together, these studies in conjunction with earlier findings on the same mouse model of mitochondrial cardiomyopathy [28], have identified several perturbed metabolites and metabolic pathways to be used in panels of biomarkers for the diagnosis or as potential therapeutic target of *CHCHD10-*related diseases.

## Supporting information

Lipidomics analysis shows a strong lipid profile changes in Chchd10S59L/+ heart

## FIGURES LEGEND

**Figure S1: Lipidomics analysis shows a strong lipid profile changes in *Chchd10*^*S59L/+*^ heart.** (A) Expression level of mRNAs coding for genes involved in carnitine and fatty acid metabolism in heart of *Chchd10*^*S59L/+*^ and wild-type mice at pre-symptomatic stage. RPLP0 is used as internal control of gene expression. GBBH: gamma-butyrobetaine dioxygenase; CPTII: carnitine-palmitoyl-transferase-2; CRAT: carnitine acetyltransferase; ACADM: Acyl-CoA deshydrogenase medium chain; FABP3: fatty acid binding protein 3 and ACSL1: Acyl-coA synthethase long chain family member 1. Data are presented as mean ± SEM compared wild type (arbitrarily set at 1), n=4-6 per group, unpaired student’s t test. (B) Number of total lipids identified and (C) number of lipids significantly affected by CHCHD10^S59L^ mutation in heart of symptomatic mice. (D) Principal component analysis (PCA) of the lipidome from *Chchd10*^*S59L/+*^ (red) and wild-type (green) heart at symptomatic stage. (E) Fraction of the total number of identified lipids in each lipid class that are significantly increased (red), significantly decreased (blue) or unchanged (grey; n.s: non-significant) in heart at the symptomatic stage. DG: diglycerides; MG: monoglyserides, LysoPC: lysophosphatidylcholine; PC: phosphatidylcholine; LysoPE: lysophosphatidylethanolamines, PE: phosphatidylethanolamine; PS: phosphatidylserine; PG: prostaglandins; SFA: saturated fatty acid; UFA: unsaturated fatty acids. (F) PCA of the metabolome from *Chchd10*^*S59L/+*^ (red) and wild-type (green) heart at symptomatic stage.

## ACKNOWLEDGEMENTS

We thank Benoit Petit-Demoulière, and Yann Herault from Phenomin (Institut Clinique de la souris, 1 rue Laurent Fries, 67404 ILLKIRCH cedex 2—CNRS, UMR7104, Illkirch, France—INSERM, U964, Illkirch, France—Université de Strasbourg, France for baseline biochemical analyses in mice and We thank Conseil General des AM et la région PACA et Corse for their financial support. This work was supported by la Ligue Nationale Contre le Cancer, by the French Government (National Research Agency, ANR) through the ‘Investments for the Future’ programs LABEX SIGNALIFE ANR-11-LABX-0028-01 and IDEX UCAJedi ANR-15-IDEX-01 by ANR ANR-16-CE16-0024-01 and by la Ville de Nice.

## REFERENCES

[1] Pfanner N, van der Laan M, Amati P, Capaldi RA, Caudy AA, Chacinska A, et al. Uniform nomenclature for the mitochondrial contact site and cristae organizing system. The Journal of cell biology. 2014;204:1083–6.

[2] Genin EC, Plutino M, Bannwarth S, Villa E, Cisneros-Barroso E, Roy M, et al. CHCHD10 mutations promote loss of mitochondrial cristae junctions with impaired mitochondrial genome maintenance and inhibition of apoptosis. EMBO Molecular Medicine. 2015;8:58–72.

[3] Burstein SR, Valsecchi F, Kawamata H, Bourens M, Zeng R, Zuberi A, et al. In vitro and in vivo studies of the ALS-FTLD protein CHCHD10 reveal novel mitochondrial topology and protein interactions. Human Molecular Genetics. 2017;27:160–77.

[4] Straub IR, Janer A, Weraarpachai W, Zinman L, Robertson J, Rogaeva E, et al. Loss of CHCHD10–CHCHD2 complexes required for respiration underlies the pathogenicity of a CHCHD10 mutation in ALS. Human Molecular Genetics. 2017;27:178–89.

[5] Anderson CJ, Bredvik K, Burstein SR, Davis C, Meadows SM, Dash J, et al. ALS/FTD mutant CHCHD10 mice reveal a tissue-specific toxic gain-of-function and mitochondrial stress response. Acta neuropathologica. 2019.

[6] Bannwarth S, Ait-El-Mkadem S, Chaussenot A, Genin EC, Lacas-Gervais S, Fragaki K, et al. A mitochondrial origin for frontotemporal dementia and amyotrophic lateral sclerosis through CHCHD10 involvement. Brain. 2014;137:2329–45.

[7] Genin EC, Madji Hounoum B, Bannwarth S, Fragaki K, Lacas-Gervais S, Mauri-Crouzet A, et al. Mitochondrial defect in muscle precedes neuromuscular junction degeneration and motor neuron death in CHCHD10(S59L/+) mouse. Acta neuropathologica. 2019.

[8] Dupuis L, Oudart H, René F, de Aguilar J-LG, Loeffler J-P. Evidence for defective energy homeostasis in amyotrophic lateral sclerosis: benefit of a high-energy diet in a transgenic mouse model. Proceedings of the National Academy of Sciences. 2004;101:11159–64.

[9] Fergani A, Oudart H, De Aguilar J-LG, Fricker B, René F, Hocquette J-F, et al. Increased peripheral lipid clearance in an animal model of amyotrophic lateral sclerosiss?. Journal of lipid research. 2007;48:1571–80.

[10] Desport J-C, Torny F, Lacoste M, Preux P-M, Couratier P. Hypermetabolism in ALS: correlations with clinical and paraclinical parameters. Neurodegenerative Diseases. 2005;2:202–7.

[11] Nakamura R, Kurihara M, Ogawa N, Kitamura A, Yamakawa I, Bamba S, et al. Investigation of the prognostic predictive value of serum lipid profiles in amyotrophic lateral sclerosis: roles of sex and hypermetabolism. Scientific Reports. 2022;12:1826.

[12] Nakamura M, Sadoshima J. Mechanisms of physiological and pathological cardiac hypertrophy. Nature Reviews Cardiology. 2018;15:387–407.

[13] Piquereau J, Caffin F, Novotova M, Lemaire C, Veksler V, Garnier A, et al. Mitochondrial dynamics in the adult cardiomyocytes: which roles for a highly specialized cell? Frontiers in physiology. 2013;4:102.

[14] Folch J, Lees M, Sloane Stanley GH. A simple method for the isolation and purification of total lipids from animal tissues. J biol Chem. 1957;226:497–509.

[15] Rozier R, Paul R, Madji Hounoum B, Villa E, Mhaidly R, Chiche J, et al. Pharmacological preconditioning protects from ischemia/reperfusion-induced apoptosis by modulating Bcl-xL expression through a ROS-dependent mechanism. The FEBS Journal. 2021;288:3547–69.

[16] Livak KJ, Schmittgen TD. Analysis of relative gene expression data using real-time quantitative PCR and the 2-ΔΔCT method. methods. 2001;25:402–8.

[17] Sud M, Fahy E, Cotter D, Azam K, Vadivelu I, Burant C, et al. Metabolomics Workbench: An international repository for metabolomics data and metadata, metabolite standards, protocols, tutorials and training, and analysis tools. Nucleic acids research. 2016;44:D463–D70.

[18] Flier JS. Starvation in the Midst of Plenty: Reflections on the History and Biology of Insulin and Leptin. Endocrine Reviews. 2018;40:1–16.

[19] Harris RB. Direct and indirect effects of leptin on adipocyte metabolism. Biochimica et Biophysica Acta (BBA)-Molecular Basis of Disease. 2014;1842:414–23.

[20] Pereira S, Cline DL, Glavas MM, Covey SD, Kieffer TJ. Tissue-specific effects of leptin on glucose and lipid metabolism. Endocrine reviews. 2021;42:1–28.

[21] Barr VA, Malide D, Zarnowski MJ, Taylor SI, Cushman SW. Insulin stimulates both leptin secretion and production by rat white adipose tissue. Endocrinology. 1997;138:4463–72.

[22] Lanznaster D, Bejan-Angoulvant T, Patin F, Andres CR, Vourc’h P, Corcia P, et al. Plasma creatinine and amyotrophic lateral sclerosis prognosis: a systematic review and meta-analysis. Amyotrophic Lateral Sclerosis and Frontotemporal Degeneration. 2019;20:199–206.

[23] Mitsumoto H, Garofalo DC, Santella RM, Sorenson EJ, Oskarsson B, Fernandes JaM, et al. Plasma creatinine and oxidative stress biomarkers in amyotrophic lateral sclerosis. Amyotrophic Lateral Sclerosis and Frontotemporal Degeneration. 2020;21:263–72.

[24] Guo QF, Hu W, Xu LQ, Luo H, Wang N, Zhang QJ. Decreased serum creatinine levels predict short survival in amyotrophic lateral sclerosis. Annals of clinical and translational neurology. 2021;8:448–55.

[25] González De Aguilar J-L. Lipid Biomarkers for Amyotrophic Lateral Sclerosis. Frontiers in Neurology. 2019;10.

[26] Blasco H, Lanznaster D, Veyrat-Durebex C, Hergesheimer R, Vourch P, Maillot F, et al. Understanding and managing metabolic dysfunction in Amyotrophic Lateral Sclerosis. Expert Review of Neurotherapeutics. 2020;20:907–19.

[27] Candelise N, Salvatori I, Scaricamazza S, Nesci V, Zenuni H, Ferri A, et al. Mechanistic Insights of Mitochondrial Dysfunction in Amyotrophic Lateral Sclerosis: An Update on a Lasting Relationship. Metabolites. 2022;12:233.

[28] Sayles NM, Southwell N, McAvoy K, Kim K, Pesini A, Anderson CJ, et al. Mutant CHCHD10 causes an extensive metabolic rewiring that precedes OXPHOS dysfunction in a murine model of mitochondrial cardiomyopathy. Cell Reports. 2022;38:110475.

[29] Lei Q, Yi T, Li H, Yan Z, Lv Z, Li G, et al. Ubiquitin C-terminal hydrolase L1 (UCHL1) regulates post-myocardial infarction cardiac fibrosis through glucose-regulated protein of 78 kDa (GRP78). Scientific Reports. 2020;10:10604.

[30] Sun H, Olson KC, Gao C, Prosdocimo DA, Zhou M, Wang Z, et al. Catabolic defect of branched-chain amino acids promotes heart failure. Circulation. 2016;133:2038–49.

[31] Kato T, Niizuma S, Inuzuka Y, Kawashima T, Okuda J, Tamaki Y, et al. Analysis of metabolic remodeling in compensated left ventricular hypertrophy and heart failure. Circulation: Heart Failure. 2010;3:420–30.

[32] Vandoorne T, De Bock K, Van Den Bosch L. Energy metabolism in ALS: an underappreciated opportunity? Acta neuropathologica. 2018;135:489–509.

[33] Paradies G, Paradies V, Ruggiero FM, Petrosillo G. Role of Cardiolipin in Mitochondrial Function and Dynamics in Health and Disease: Molecular and Pharmacological Aspects. Cells. 2019;8.

[34] Sol J, Jové M, Povedano M, Sproviero W, Domínguez R, Piñol-Ripoll G, et al. Lipidomic traits of plasma and cerebrospinal fluid in amyotrophic lateral sclerosis correlate with disease progression. Brain Communications. 2021;3.

[35] Pettegrew J, Levine J, McClure R. Acetyl-L-carnitine physical-chemical, metabolic, and therapeutic properties: relevance for its mode of action in Alzheimer’s disease and geriatric depression. Molecular psychiatry. 2000;5:616–32.

[36] Kerner J, Hoppel C. Fatty acid import into mitochondria. Biochimica et Biophysica Acta (BBA)-Molecular and Cell Biology of Lipids. 2000;1486:1–17.

[37] van Vlies N, Ferdinandusse S, Turkenburg M, Wanders RJ, Vaz FM. PPARα-activation results in enhanced carnitine biosynthesis and OCTN2-mediated hepatic carnitine accumulation. Biochimica et Biophysica Acta (BBA)-Bioenergetics. 2007;1767:1134–42.

[38] Stanley WC, Recchia FA, Lopaschuk GD. Myocardial Substrate Metabolism in the Normal and Failing Heart. Physiological Reviews. 2005;85:1093–129.

[39] Lopaschuk GD, Jaswal JS. Energy metabolic phenotype of the cardiomyocyte during development, differentiation, and postnatal maturation. Journal of cardiovascular pharmacology. 2010;56:130–40.

[40] Calabrese V, Ravagna A, Colombrita C, Scapagnini G, Guagliano E, Calvani M, et al. Acetylcarnitine induces heme oxygenase in rat astrocytes and protects against oxidative stress: involvement of the transcription factor Nrf2. Journal of neuroscience research. 2005;79:509–21.

[41] Patel SP, Sullivan PG, Lyttle TS, Rabchevsky AG. Acetyl-l-carnitine ameliorates mitochondrial dysfunction following contusion spinal cord injury. Journal of Neurochemistry. 2010;114:291–301.

[42] Di Cesare Mannelli L, Ghelardini C, Calvani M, Nicolai R, Mosconi L, Vivoli E, et al. Protective effect of acetyl-L-carnitine on the apoptotic pathway of peripheral neuropathy. European Journal of Neuroscience. 2007;26:820–7.

[43] Makrecka-Kuka M, Sevostjanovs E, Vilks K, Volska K, Antone U, Kuka J, et al. Plasma acylcarnitine concentrations reflect the acylcarnitine profile in cardiac tissues. Scientific Reports. 2017;7:17528.

[44] Strand E, Pedersen ER, Svingen GFT, Olsen T, Bjørndal B, Karlsson T, et al. Serum Acylcarnitines and Risk of Cardiovascular Death and Acute Myocardial Infarction in Patients With Stable Angina Pectoris. Journal of the American Heart Association. 2017;6:e003620.

