## Supplementary figures and images for "Multiomics study of *CHCHD10^S59L^*-related disease reveals energy metabolism downregulation: OXPHOS and β-oxidation deficiencies associated with lipids alterations"

### Lipidomics analysis shows a strong lipid profile changes in Chchd10S59L/+ heart

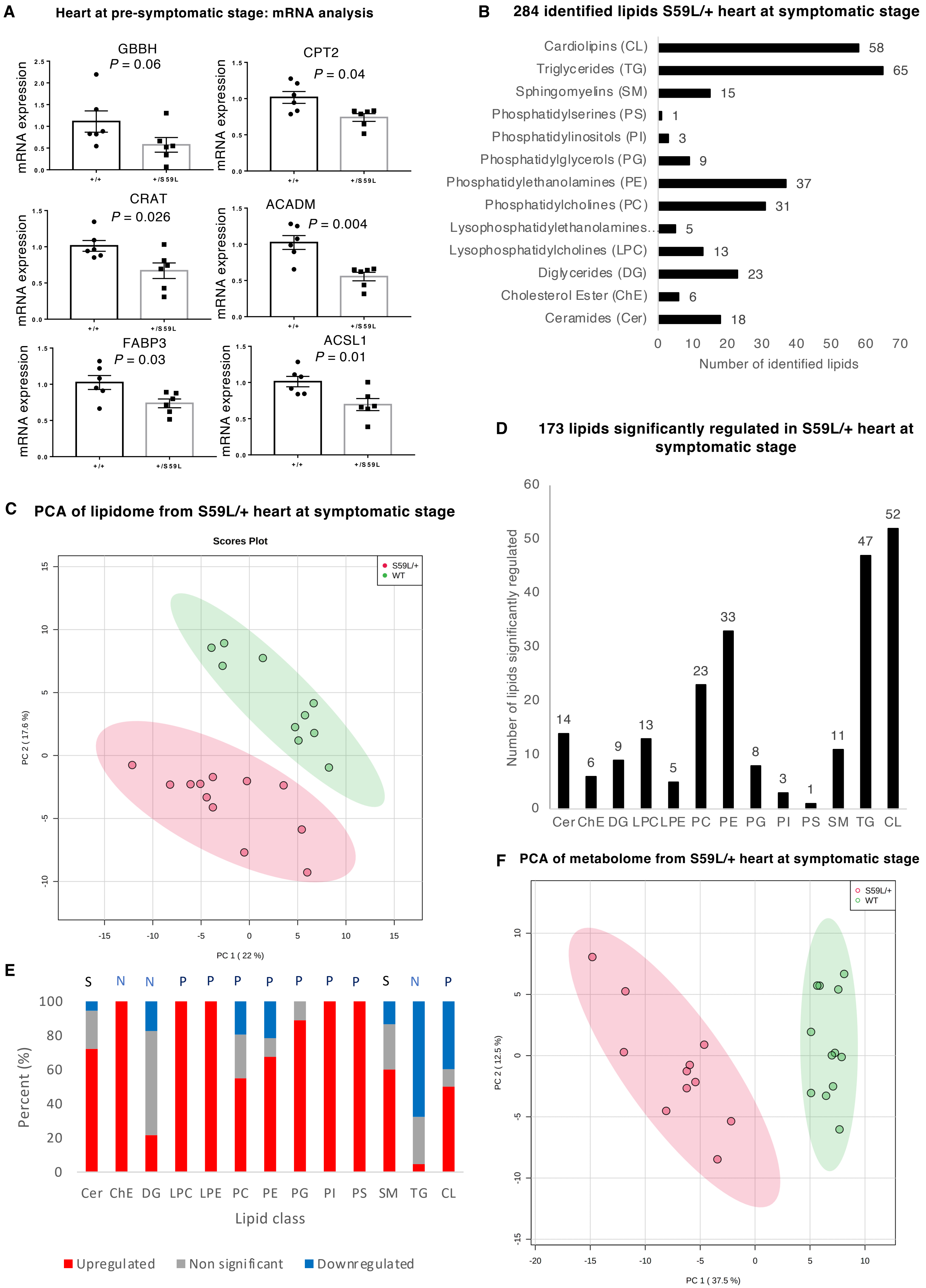
