## Supplementary material for "Multiomics study of *CHCHD10^S59L^*-related disease reveals energy metabolism downregulation: OXPHOS and β-oxidation deficiencies associated with lipids alterations": supplmentary material and methods

| **Gene** | **Forward sequences** | **Reverse sequences** |
| --- | --- | --- |
| CRAT | AGAGCCTGTTGGCATCCTAACC | TTGTCCAGGCACACGGTGAAGA |
| CPT2 | GATGGCTGAGTGCTCCAAATACC | GCTGCCAGATACCGTAGAGCAA |
| GBBH | AGCTTCCACACTGACTACCCAG | CTCCTTGAGCTTTTGGCACACG |
| FABP3 | AGAGTTCGACGAGGTGACAGCA | TTGTCTCCTGCCCGTTCCACTT |
| ACADM | AGGATGACGGAGCAGCCAATGA | TTGTCTCCTGCCCGTTCCACTT |
| ACSL1 | ATCAGGCTGCTTATGGACGACC | CCAACAGCCATCGCTTCAAGGA |

**Table S1**: Primers for RT-qPCR analysis used in this study

| **Antigen** | **Source** | **Host** | **Dilution** |
| --- | --- | --- | --- |
| PHGDH | Protein Tech (# 14719-1-AP) | Rabbit | 1/2000 |
| PSAT1 | Protein Tech (# 10501-1-AP) | Rabbit | 1/2000 |
| ASNS | Protein Tech (# 10501-1-AP) | Rabbit | 1/2000 |
| PSPH | Protein Tech (# 14513-1-AP) | Rabbit | 1/1000 |
| SHMT2 | Cell signalling (# 12762) | Rabbit | 1/1000 |
| ERK2 | Santa-Cruz Biotechnology (# sc-1647) | Mouse | 1/1000 |
| Rabbit HRP | Cell signalling (# 7074S) | Goat | 1/5000 |
| Mouse HRP | Cell signalling (# 7076S) | Horse | 1/3000 |

**Table S2**: Antibodies used for western blot in this study

| **Metabolites** | **VIP** |
| --- | --- |
| PAF-C16 | 3.5068 |
| LPC(18:0) | 3.2951 |
| LPC(16:0) | 3.2508 |
| LPC(20:3) | 3.1861 |
| PC(O-14:0/2:0) | 3.1149 |
| PC(O-18:2/2:0) | 3.0054 |
| LPC(20:2) | 2.974 |
| N-ACETYLGLYCINE | 2.9379 |
| LPC(14:0) | 2.879 |
| LPC(18:1) | 2.8552 |
| PC(38:2) | 2.8518 |
| 3-AMINO-4-HYDROXYBENZOIC_ACID | 2.8042 |
| LPC(22:4) | 2.7155 |
| PC(O-20:4/2:0) | 2.6802 |
| PAF-C18 | 2.6781 |
| PC(O-16:1/2:0) | 2.6596 |
| PC(36:3) | 2.6331 |
| LPC(20:0) | 2.6061 |
| PC(33:0) | 2.5694 |
| LPC(16:1) | 2.5261 |
| LPC(22:1) | 2.5174 |
| PC(34:1) | 2.4876 |
| PC(36:1) | 2.4604 |
| PC(34:3) | 2.4604 |
| PC(25:0) | 2.4589 |
| SM(40:2) | 2.4521 |
| PC(32:1) | 2.4289 |
| L-ASPARTATE | 2.4148 |
| PALMITATE | 2.4094 |
| PC(O-16:2/2:0) | 2.4083 |
| PC(36:2) | 2.3994 |
| GAMMA-LINOLENIC_ACID | 2.382 |
| LPC(18:2) | 2.3687 |
| PC(40:7) | 2.337 |
| ALPHA-GLUCOSE | 2.331 |
| GLUCURONIC_ACID | 2.3236 |
| SM(40:3) | 2.2867 |
| PC(38:5) | 2.2575 |
| L-GLUTAMIC_ACID | 2.24 |
| PC(O-20:6/2:0) | 2.2378 |
| PC(32:2) | 2.2362 |
| PC(O-14:1/2:0) | 2.2265 |
| RIBOSE_5-PHOSPHATE | 2.2158 |
| PC(30:0) | 2.2053 |
| 3-SULFINO-L-ALANINE | 2.1819 |
| PC(40:6) | 2.164 |
| PC(38:6) | 2.1458 |
| LPC(20:4) | 2.1424 |
| PC(36:5) | 2.1419 |
| LPE(18:2) | 2.1257 |
| SM(32:2) | 2.0193 |
| LPC(22:6) | 2.0155 |
| ADENOSINE_5'-MONOPHOSPHATE | 2.0031 |
| STEARATE | 1.987 |
| PC(35:3) | 1.9832 |
| GLUCONIC_ACID | 1.9696 |
| MANNITOL | 1.949 |
| PC(33:2) | 1.8787 |
| PC(35:1) | 1.8136 |
| PC(21:0) | 1.8037 |
| PC(O-18:4/2:0) | 1.7804 |
| PC(31:0) | 1.7467 |
| PC(36:4) | 1.713 |
| PC(O-12:0/2:0) | 1.7014 |
| N-ACETYLNEURAMINATE | 1.701 |
| C14-CARNITINE | 1.6783 |
| LPE(16:0) | 1.6352 |
| PC(34:2) | 1.6216 |
| GLUCOSE_6-PHOSPHATE | 1.5737 |
| PC(35:2) | 1.511 |
| PC(8:0) | 1.4925 |
| PC(37:4) | 1.4659 |
| GUANOSINE_5'-MONOPHOSPHATE | 1.4484 |
| GLYCERATE | 1.4207 |
| 3-DEHYDROSHIKIMATE | 1.3884 |
| ACETYLCHOLINE | 1.3644 |
| XANTHINE | 1.3116 |
| PC(32:0) | 1.3013 |
| LINOLEATE | 1.257 |
| ELAIDIC_ACID | 1.195 |
| MALONATE | 1.1722 |
| PC(O-16:3/2:0) | 1.1438 |
| LAUROYLCARNITINE | 1.1156 |
| CHOLESTERYL_ACETATE | 1.0935 |
| NICOTINAMIDE | 1.0407 |
| ALPHA-AMINOADIPATE | 1.0261 |

**Table S3**: Detailed information of Metabolites identified in plasma by differential analysis (VIP score > 1) in OPLS-DA model.

| **Metabolites identified in plasma from pre-symptomatic mice** | **FC (KI/WT)** | ***P* Value** |
| --- | --- | --- |
| **INOSINE_5'-MONOPHOSPHATE** | **0.1166** | **0.02** |
| **4-IMIDAZOLEACETIC_ACID** | **1.9784** | **0.02** |
| **N-ACETYL-L-PHENYLALANINE** | **1.9699** | **0.02** |
| **N-ACETYLPUTRESCINE** | **1.819** | **0.02** |
| **CETOGLUTARATE** | **2.9576** | **0.03** |
| **L-LYSINE** | **3.3578** | **0.03** |
| **TRIGONELLINE** | **2.3121** | **0.03** |
| **PANTOTHENIC_ACID** | **2.0064** | **0.03** |
| **N-ACETYLGLYCINE** | **0.54273** | **0.03** |
| **2-HYDROXY-4-(METHYLTHIO) BUTYRIC_ACID** | **1.5835** | **0.03** |
| **ALLANTOIN** | **1.5466** | **0.03** |

**Table S4**: Metabolites significantly changed in plasma from Chchd10^S59L^ mice compared to wild-type at the pre-symptomatic stage. Fold change, and p value < 0.05 (by the Mann- Whitney U test).

| **Metabolites identified in plasma from symptomatic mice** | **FC (KI/WT)** | ***P* Value** |
| --- | --- | --- |
| **BETAINE** | **1.152** | **0.001** |
| **PANTOTHENIC_ACID** | **2.3786** | **0.002** |
| **C3-CARNITINE** | **2.6869** | **0.005** |
| **L-VALINE** | **1.5925** | **0.005** |
| **PAF-C16** | **0.65757** | **0.006** |
| **L-ALANINE** | **2.548** | **0.008** |
| **GUANIDINOACETATE** | **2.1889** | **0.008** |
| **DEOXYCARNITINE** | **2.0146** | **0.008** |
| **L-METHIONINE** | **1.8298** | **0.008** |
| **4-AMINOBUTANOATE** | **1.8236** | **0.008** |
| **LEUCINE_NORLEUCINE** | **1.5586** | **0.008** |
| **L-PROLINE** | **2.0444** | **0.012** |
| **HISTAMINE** | **2.4714** | **0.014** |
| **TAURINE** | **1.733** | **0.014** |
| **LPC(16:1)** | **0.6304** | **0.014** |
| **PC(36:2)** | **0.72354** | **0.014** |
| **PC(O-16:1/2:0)** | **0.74765** | **0.014** |
| **CHOLESTERYL_ACETATE** | **0.71699** | **0.018** |
| **L-THREONINE** | **2.0657** | **0.018** |
| **LPC(20:0)** | **0.40223** | **0.022** |
| **L-SERINE** | **1.9736** | **0.022** |
| **ADENINE** | **1.9626** | **0.022** |
| **O-ACETYL-L-CARNITINE** | **1.5831** | **0.022** |
| **METHYLGUANIDINE** | **1.4234** | **0.022** |
| **3-AMINO-4-HYDROXYBENZOIC_ACID** | **0.72766** | **0.022** |
| **LPC(18:1)** | **0.73607** | **0.022** |
| **ALPHA-GLUCOSE** | **0.58194** | **0.027** |
| **C2-CARNITINE** | **1.5897** | **0.027** |
| **L-ASPARTATE** | **0.38855** | **0.032** |
| **CREATINE** | **1.7434** | **0.032** |
| **CYTIDINE_2'** | **2.154** | **0.035** |
| **ADENOSINE** | **2.9784** | **0.035** |
| **PAF-C18** | **0.34532** | **0.035** |
| **C4-CARNITINE** | **2.3422** | **0.035** |
| **FUMARATE** | **2.2806** | **0.035** |
| **4-HYDROXY-L-PROLINE** | **2.2579** | **0.035** |
| **CITRULLINE** | **1.9757** | **0.035** |
| **BETA-ALANINE** | **1.7197** | **0.035** |
| **3-SULFINO-L-ALANINE** | **0.58242** | **0.035** |
| **L-TRYPTOPHAN** | **1.6439** | **0.035** |
| **LPC(18:0)** | **0.6475** | **0.035** |
| **LPC(14:0)** | **0.69051** | **0.035** |
| **PC(O-14:0/2:0)** | **0.69876** | **0.035** |
| **PC(36:1)** | **0.73634** | **0.035** |
| **L-TYROSINE** | **1.5574** | **0.038** |
| **LPC(16:0)** | **0.73816** | **0.038** |
| **LPC(22:1)** | **0.53134** | **0.038** |
| **XANTHINE** | **0.41168** | **0.045** |
| **PC(32:1)** | **0.68128** | **0.045** |

**Table S5**: Metabolites significantly changed in plasma from Chchd10^S59L^ mice compared to wild-type at the symptomatic stage. Fold change, and p value < 0.05 (by the Mann- Whitney U test).
